# Estimating driver-tissues by robust selective expression of genes associated with complex diseases or traits

**DOI:** 10.1101/491878

**Authors:** Lin Jiang, Chao Xue, Shangzhen Chen, Sheng Dai, Peikai Chen, Pak Chung Sham, Haijun Wang, Miaoxin Li

## Abstract

The driver tissues or cell-types of many human diseases, in which susceptibility genes cause the diseases, remain elusive. We developed a framework to detect the causal-tissues of complex diseases or traits according to selective expression of disease-associated genes in genome-wide association study (GWAS). The core method of the framework is a new robust z-score to estimate genes’ expression selectivity. Through extensive computing simulations and comparative analyses in a large-scale schizophrenia GWAS, we demonstrate the robust z-score is more sensitive than existing methods to detect multiple selectively expressed tissues, which further lead to the estimation of more biological sensible driver tissues. The effectiveness of this framework is further validated in five representative complex diseases with the usage of GWAS summary statistics and transcript-level expression in GTEx project. Finally, we also demonstrate that the prioritized tissues and the robust selective expression can enhance characterization of directly associated genes of a disease as well. Interesting results include the estimation of lung as a driver tissue of rheumatoid arthritis, consistent with clinical observations of morbidity between rheumatoid arthritis and lung diseases.

## Introduction

Tissue-selectivity is an important feature of many complex diseases [1] and the selective expression of their susceptibility genes may determine the tissue-selective pathology [2, 3]. It is common that a complex disease often involves multiple affected tissues; and it is usually tricky to characterize the causal or driver tissues [4]. This can be explained by an example of schizophrenia. It is certain that brain must be a relevant organ of schizophrenia. However, as human brains consist of multiple heterogeneous areas, it is crucial to know which areas are the exact drivers [5]. Besides, our current knowledge on the tissue selectivity of complex diseases is often biased toward clinical observations. It should be noted many clinically observed tissues are not necessarily the driver- or causal tissues. For most of human diseases, the primary driver tissues remain elusive [6]. Many studies showed that disease causal genes tend to have elevated selective expression in the pathogenic tissues [1, 2], implicating a basis for the tissue selectivity or driver of diseases. Analyses of genes’ selective expression profiles can expand the knowledge on human diseases[7] and even facilitate characterizing new causal genes [8]. Ongen et al. proposed to estimate the causal tissues for complex traits and diseases by measuring the GWAS-associated variants’ eQTL activity in different tissues[9]. But this study did not straightforwardly consider genes’ selectivity expression and its requirement of eQTL may also limit the application of their approach.

Tissue selective expression refers to much higher or lower expression of a gene in one or some minority tissues than majority tissues[10]. However, it is difficult to quantify the relative difference due to ambiguous boundaries between the minority and the majority in practice. There are several methods for detecting tissue selective expression of genes (See method description in the review [11]). Most early methods can only tell whether a gene has overall selective expression [12, 13]; and most recent methods are underpowered to detect selective expression at individual tissues when there is more than one tissue with selective expression [14]. Several large collaborative projects including GTEx [15-17] [18, 19] have produced transcriptomes across many human tissues. These ever-increasing resources are calling for more powerful selective expression measures and more studies on tissue-selective pathology of human diseases.

In this study we proposed a framework to estimate <u>d</u>river-tissues or cell-types of complex diseases or traits by robust <u>s</u>elective <u>e</u>xpression (DIRSE) of genes derived from genome-wide association study (GWAS) summary statistics. The core method of the framework is a novel robust z-score for detecting selectively expressed tissues. After examining the statistical performance of the robust z-score, we applied it to produce selective expression profiles at transcript- and gene-levels in 50 tissues and cell-types from GTEx Project [18]. Using summary statistics from a schizophrenia GWAS, we compared its effectiveness of identifying schizophrenia’s driver tissues with three alternative selective expression measures. We further applied this approach to identify potential driver-tissues of 5 representative complex diseases and investigated how the prioritized tissues can enhance detection of susceptibility genes in post GWAS analyses. For simplicity, we omit the terminology cell-types throughout the paper, but it should be noted the statement of tissues also includes cell-types in the present paper.

## Results

### The proposed framework of estimating driver tissues and its robust z-score for selective expression

The framework aims to estimate driver tissues by tissue-selective expression of GWAS genes (See the workflow in Figure 1). The assumption is that the tissue selective expression of causal or susceptibility genes determines the tissues where complex diseases happen primarily, which are called driver tissues. Therefore, a cause tissue is very likely to have an enrichment of selective expression by the susceptibility genes of a disease. The inputs of the framework include two types of data, standardized expression values of multiple tissues and GWAS summary statistics or p-values for a certain disease at variants. The expression values at genes and transcripts are used to evaluate selective expression by a novel robust score. The GWAS p-values are used to detect susceptibility genes by a conditional gene-based association test we published recently [20]. The driver tissues are estimated by enrichment analysis of GWAS associated genes with selective expression. The selective expression and enrichment analyses are implemented in an online tool, http://grass.cgs.hku.hk:8080/dirse/.

**Figure 1:**
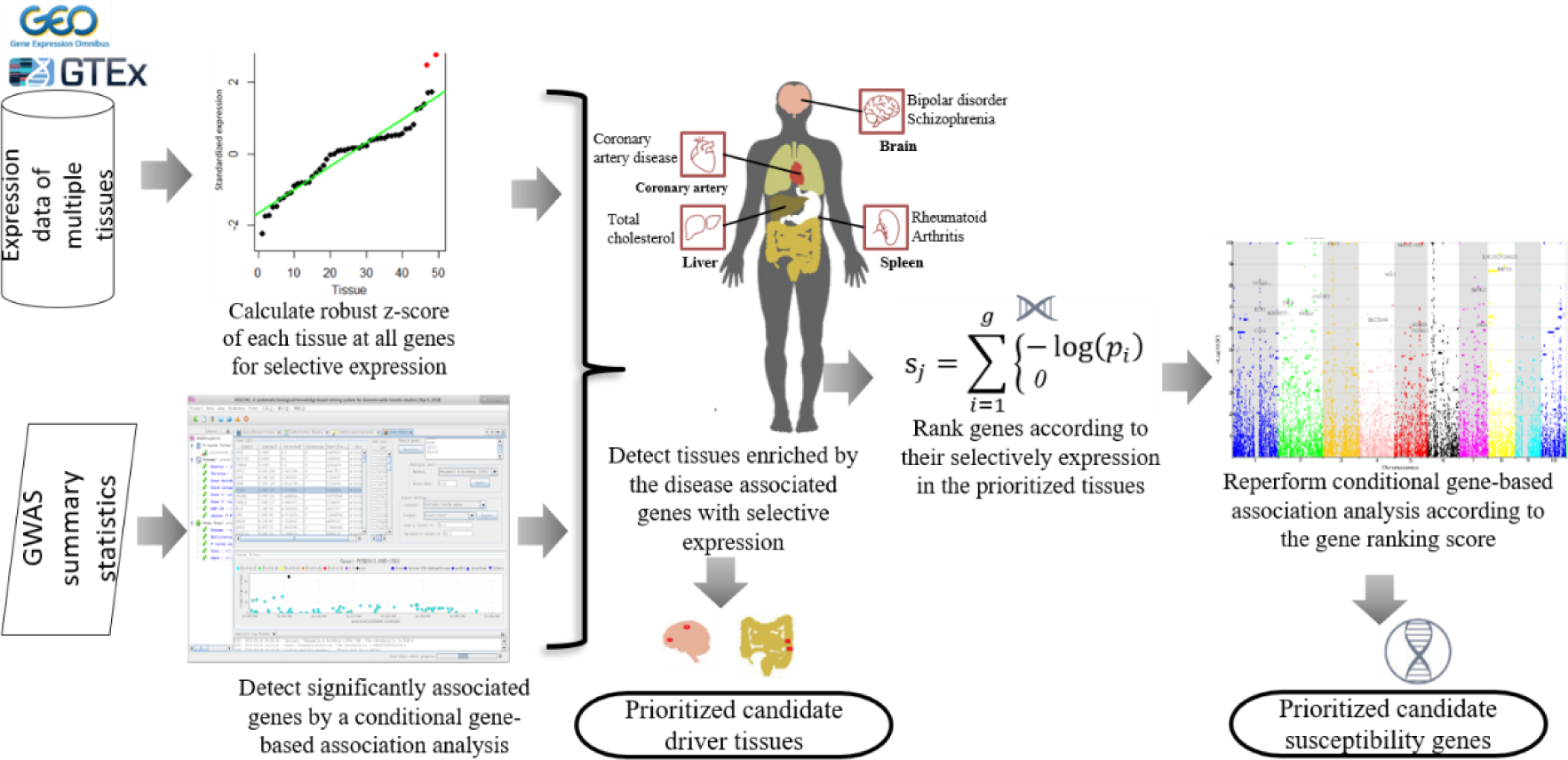
Analysis procedure to prioritize tissue selectivity and susceptibility genes of complex diseases. The expression of multiple tissues (from GTEx) is input to measure selective expression of tissues by a robust z-score. The GWAS p-values are used to detect genes associated with a complex disease by a conditional gene-based association test on KGG. The causal tissues are then estimated by enrichment analysis of GWAS-genes with selective expression. The selective expression in the causal tissues is used to prioritize susceptibility genes by the conditional gene-based association test in turn.

The core method of the framework is the robust z-score for detecting tissue selective expression. It subtly constructs a fitted line for ranked expression at a gene, and integrates expression deviation and expression variation to measure a robust selective expression (See details in the Method section). Under null hypothesis, it can approximate valid p-values (See the QQ plots in Figure S2 and S3), which will greatly facilitate statistical inference of selective expression. Extensive computer simulations show that the robust z-score is more powerful than the conventional z-score when there are multiple selectively expressed tissues (Table S1). This advantage lends itself to public datasets (including GTEx) in which the number of selectively expressed tissues and sample sizes of each tissue are often variable.

### Tissue selective expression profiles in 50 tissues produced by the robust z-score

The robust z-score approach was then applied to generate tissue selective expression profiles by using RNA-Seq data from GTEx project (V7) [18] after stringent quality control (See details in Methods and Material Sections). While the number of selectively expressed genes varies from tissues to tissues (Table S3), the profiles have three common interesting features. First, the pair-wise correlations based on the robust z-scores are substantially different from those based on the original expression values (TPM). Many tissues have the correlation over 0.7 (Pearson coefficients) based on the expression values (Figure S4 and S5) while most of them have zero correlation based on the robust z-score (Figure S6 and S7). However, tissues of similar origins have high Pearson correlations (>0.8) based on the robust z-scores, such as Brain-Spinalcord vs. Brain-Substantianigra pair, Cervix-Ectocervix vs. Vagina pair, and Skin-SunExposed(Lowerleg) vs. Skin-NotSunExposed(Suprapubic) pair. The biologically sensible consistency suggests the robust z-score scores quantify selective expression of the tissues correctly. Second, the original expression values have low correlation with the selective expression values at most genes. As shown in Figure S8 and S9, only 12 tissues have moderate Spearman correlation co-efficient, *r*∈[0.3,0.6] between the expression values and robust z-score values at gene levels. Most tissues have nearly zero or even negative Spearman correlations. For example, the left heart ventricle has a negative correlation, −0.21. In the left heart ventricle, the MT-CO1 gene has the largest expression value, 77005.9 TPM while the robust z-score is only 3.3. This is also true for many other mitochondrial protein coding genes, e.g., MT-ATP6, MT-CO3 and MT-ND4. On the other hand, some genes have low or moderate expression values but very high robust z-scores, e.g., LINC02500 (2.0 TPM vs. 912.0) and XIRP1 (251.9 TPM vs. 2392.1). Many genes encoding myosin light chain (e.g. MYL2 and MYL2) have both high expression values and large robust z-scores. The myosin light chains are critical for functions of heart cells [21] and quite a few genes encoding myosin light chains have been linked to cardiovascular diseases, e.g. MYL2[22] and MYL4 [23]. The correlation patterns of selective expression based on the transcript-level expression are similar to those based on gene levels.

Third, the expression data at transcript level lead to discovery of much more selectively expressed genes. In most tissues, the usage of transcript-level expression detects on average 54% extra selectively expressed genes which are missed by the usage of gene-level expression (Table S3). Although the conservative Bonferroni correction for multiple transcripts of a gene may lead to the missing of some selectively expressed genes, the unique genes selectively expressed according to transcript-level expression are still on average 5.5 times more than that according to gene-level expression in the 50 tested tissues. Here is an individual example. The transcript ENST00000568186 of TCF4 has very high standardized tissue selective expression score 14.8 (*p*=1.46×10 ^-49^) in brain frontal cortex while averaged expression value of the gene has no selective expression (z=0.96, *p*=0.34) in the same tissue. TCF4 is an important gene linked to multiple brain disorders including epilepsy [24], intellectual disability [25], and schizophrenia [26]. Biologically, the transcript level expression would be closer to what happens.

### Compare the performance of different selective expression methods for estimating driver tissues of schizophrenia

With the usage of expression data, we investigated the performance of the robust z-scores and three other metrics for prioritizing driver-tissues by GWAS-genes on the above proposed framework. In a large-scale Meta-GWAS study [27], 108 genes associated with schizophrenia detected by the conditional gene-based association analysis[20] were used as the benchmark dataset (See more in the method section and gene list in Excel Table S1). Table 1 shows the top five tissues enriched by the associated genes with selective expression detected by different approaches. Based on the robust selective expression by the proposed z-score, all the estimated top five driver-tissues of schizophrenia in brain. Four estimated driver tissues (anterior cingulate cortex, frontal cortex, amygdala and hippocampus) out of the top five tissues based on the gene-level selective expression are overlapped with that based on the transcript-level selective expression, suggesting the robustness of the approach. There have been numerous studies implicating them as primary causal tissues of schizophrenia e.g.,[28] [29-32]. However, the transcript-level selective expression leads to much higher significance and more selectively expressed genes than the gene-level selective expression at the estimate tissues (e.g., *p*=1.6×1 0^-6^ vs. 1.4×10 ^-9^ and n=27 vs. 44 at anterior cingulate cortex), suggesting a higher power of the former. The top tissue according to the transcript-level selective expression is the cerebellum. This supports a recent finding that the ‘little brain’ plays a major role in schizophrenia[33].

**Table 1:**
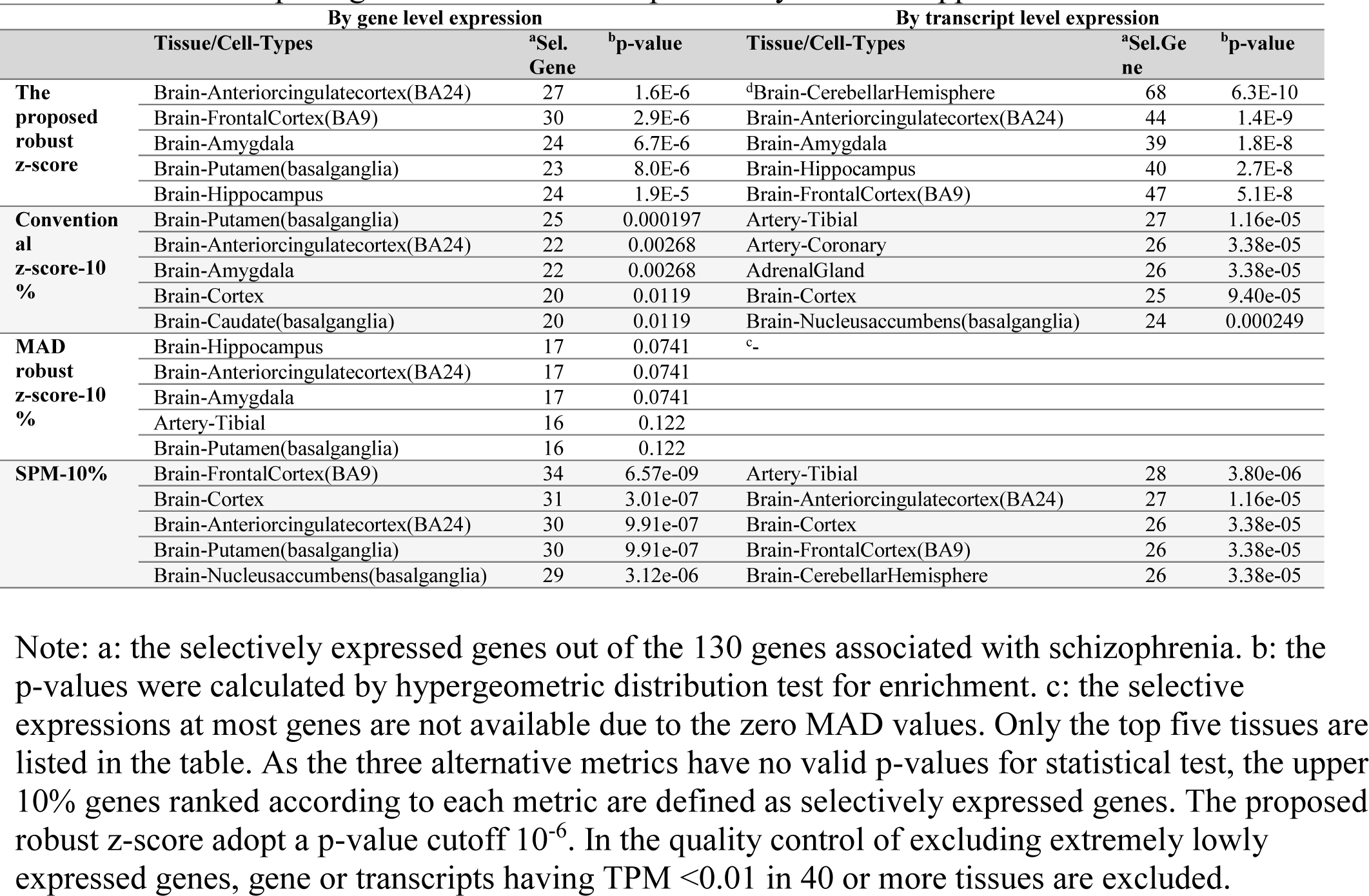
Prioritized pathogenic tissues of schizophrenia by different approaches

The estimated driver tissues according to the alternative selective expression metrics are generally less biologically sensible. The usage of conventional z-scores led to much fewer selectively expressed genes. Using the gene-level expression, it results in only one significant tissue, brain putamen. Using transcript-level expression, it ranks artery-tibial, artery-coronary and adrenal gland as the top three driver tissue of schizophrenia. There are few literatures linking these tissues to schizophrenia. Another widely-used robust z-score by median absolute deviation (MDA) with the usage of gene level expression failed to detect any significant tissues. The transcript level selective expression produced no results because of the zero MAD values. The measure of SPM [14] also prioritized many brain regions with the usage of gene-level expression. Its top five tissues belong to the brain. However, the top tissue, tibial artery, according to the transcript-level selective expression is an unlikely relevant tissue. The comparison showed the selective expression produced by proposed robust z-score is more effective than that by other metrices to prioritize driver tissue candidates of complex diseases, which is particularly true for transcript-level expression. We therefore employed the transcript-level selective expression by the proposed robust z-score to investigate tissue-selective pathogens of other complex diseases in the paper.

### Contribution of genes with low expression to the prioritization of disease related tissues

We also noticed that lowly expressed genes cannot be ignored in the prioritization of disease-related tissues. A removal of lowly expressed genes led to substantial decrease in enrichment significance when the transcript-level selective expression was used for the estimation. Table 2 shows the enrichment significance of three important brain regions for schizophrenia (cerebellar hemisphere, anterior cingulate cortex and frontal cortex) when lowly expressed genes are removed according to different cutoffs. For example, the p-values for enrichment at cerebellar hemisphere is 6.3×10 ^-10^ when genes with transcript level expression as low as 0.01 TPM is included. However, the p-value becomes 8.4×10 ^-5^ after excluding genes with transcript level expression <1 TPM at all tissues. The number of selectively expressed GWAS genes also decreases from 68 to 54 accordingly. In the other two tissues, the influence of expression cutoffs is similar. Here are some examples of individual genes. For example, CACNA1C and TCF4 are well-known candidate susceptibility genes of schizophrenia [34, 35]. CACNA1C encodes calcium voltage-gated channel subunit alpha1 C which is important for brain functions. This gene has 26 transcripts with expression in GTEx dataset and 23 has no significant expression. The transcript (ENST00000399641) has the largest selective expression z-score 210.3 in frontal cortex (*p*<1.0×10 ^-200^). However, it has only 0.2 TMP expression in frontal cortex and nearly zero TPM in most tissues. TCF4 encodes the basic Helix-Loop-Helix (bHLH) transcription factor 4, which is among the few schizophrenia-risk genes that have been consistently replicated. It has 43 transcripts with expression in GTEx dataset but only two has significant selective expression in frontal cortex. The transcript ENST00000568186 has the highest expression 0.298 TPM in frontal cortex and nearly zero TPM in most majority of tissues, which leads to a robust z-score 14 (*p*=1.6×10 ^-44^) in frontal cortex. A TPM cutoff even as low as 0.4 will exclude these important candidate genes of schizophrenia.

**Table 2:** The number of selectively expressed genes and enrichment statistical significance for different minimal expression values

|  | Cerebellar Hemisphere |  |  |  | Anterior cingulate Cortex |  |  |  | Frontal Cortex |  |  |  |
| --- | --- | --- | --- | --- | --- | --- | --- | --- | --- | --- | --- | --- |
|  | Gene-level expression |  | Transcript-level expression |  | Gene-level expression |  | Transcript-level expression |  | Gene-level expression |  | Transcript-level expression |  |
| <sup>a</sup>Cutoff | <sup>b</sup>#Gene | <sup>c</sup>p-value | #Gene | p-value | #Gene | p-value | #Gene | p-value | #Gene | p-value | #Gene | p-value |
| **0.01** | 35 | 7.9E-5 | 68 | 6.3E-10 | 27 | 1.7E-6 | 44 | 1.4E-9 | 30 | 3.0E-6 | 47 | 5.1E-8 |
| **0.05** | 35 | 8.5E-5 | 68 | 1.4E-9 | 27 | 2.8E-6 | 43 | 8.1E-9 | 30 | 4.6E-6 | 46 | 2.5E-7 |
| **0.1** | 35 | 9.2E-5 | 68 | 2.0E-9 | 27 | 4.6E-6 | 43 | 1.2E-8 | 30 | 7.7E-6 | 46 | 3.7E-7 |
| **0.5** | 35 | 1.0E-4 | 61 | 4.9E-7 | 28 | 4.9E-6 | 39 | 7.4E-7 | 31 | 9.2E-6 | 41 | 2.5E-5 |
| **1** | 35 | 6.5E-5 | 54 | 8.5E-5 | 27 | 1.5E-5 | 38 | 1.5E-6 | 31 | 9.4E-6 | 40 | 3.3E-5 |

Note: a: remove genes with expression $< c$ in all tissues. b: the number of significant genes; c: the p-value for enrichment analysis calculated by hypergeometric test. According to a cutoff x, gene or transcripts having TPM $< x$ in 40 or more tissues are excluded.

### Identify driver tissues in five representative complex diseases/traits

The above success in the benchmark GWAS dataset, schizophrenia, motivated us to validate the effectiveness of selective expression in five representative complex diseases or traits as proof-of-principle examples. Figure 2 and S10 shows the prioritized tissues. Generally, for all the five diseases/traits, not only did the proposed framework confirm previous findings, it also detected multiple new candidate driver tissues. Some of the tissue are not intuitive (e.g., lung for rheumatoid arthritis and nerve cells for coronary artery disease), which requires further experimental validation.

**Figure 2:**
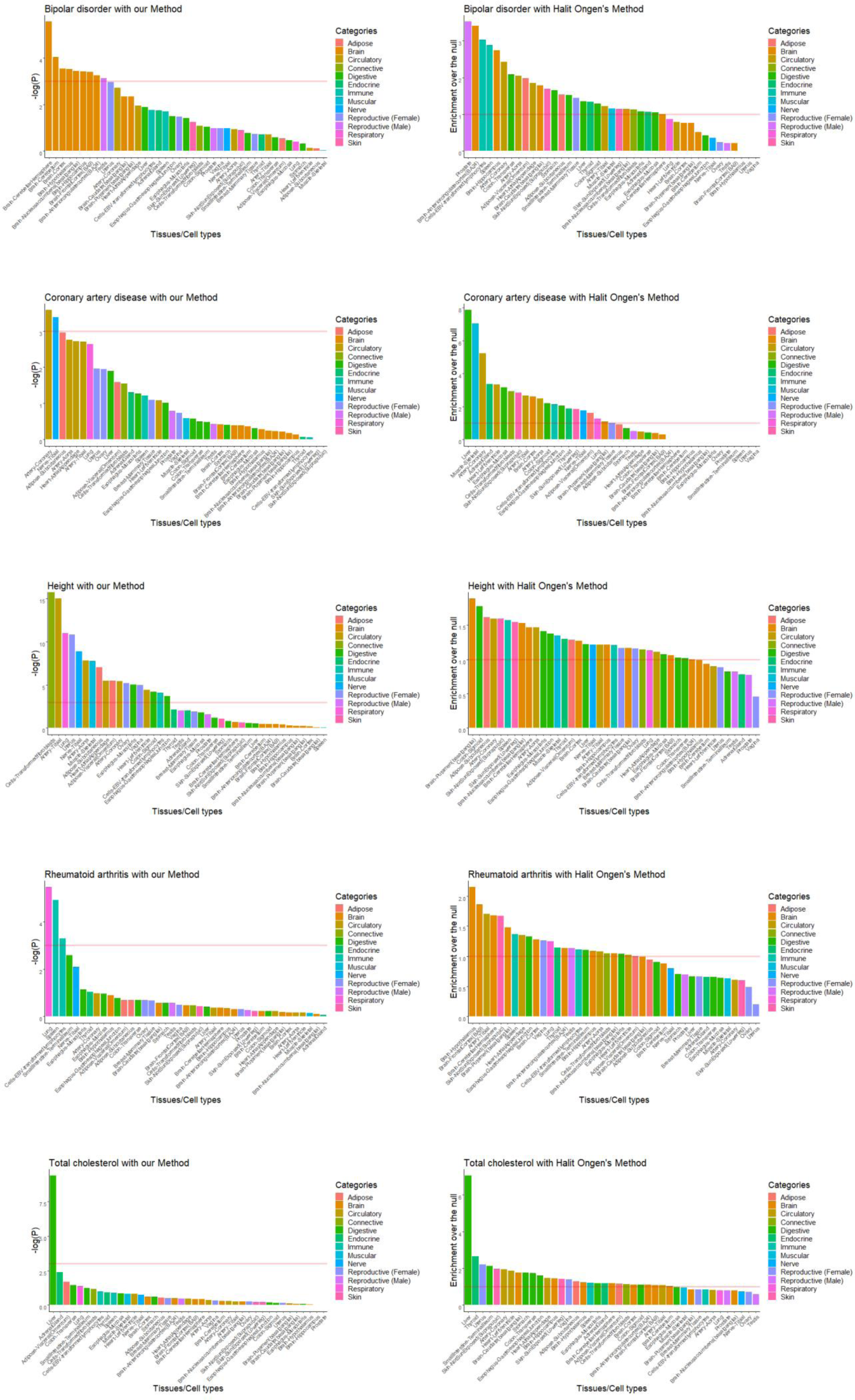
Driver tissues inferred by our proposed method and Ongen’s method in five representative complex diseases. Each row shows one disease/trait. The left part shows results inferred by our proposed method, right shows by Ongen’s method. On the *x* axis, the 41 overlapped tissues are classified into 13 groups based on anatomy, which are filled with different colors as the legend shows. The tissues are sorted by the corresponding value on *y* axis in descending order. For our proposed method, the bar on *y* axis shows the negative log_10_-transformed enrichment p value of each tissue and the red horizontal line denotes the p value cutoff by Bonferroni correction. While for Ongen’s method, the bar on the *y* axis shows the enrichment over the null of each tissue and the red horizontal line shows enrichment over the null is 1.

#### Bipolar disorder (BD)

The top ten driver tissues of bipolar disorder estimated by the proposed approach all fall into brain areas. Similar to schizophrenia, the most significant tissue is also cerebellar hemisphere (*p* =2.72×10 ^-6^), which has been implicated with BD by many studies [36]. However, it also has different prioritization results to schizophrenia. For example, brain amygdala is prioritized as the 14^th^ significant tissue for BD, while the fourth for schizophrenia, probably implying the different pathogens of the two brain disorders. Most of these significant brain areas have been suggested as pathogenic tissues of bipolar disorder, including hippocampus [37], hypothalamus [38], basal ganglia [39]. Interestingly, the brain cervical spinal cord C1 is prioritized as the second most significant driver tissue, yet there have been no studies implicating this tissue for BD by far. As a bridge between brain and other body parts, cervical spinal cord may have a potential contribution to the development of BD. This is subject to validation by experiments in the feature.

#### Coronary artery disease (CAD)

CAD causes impaired blood flow in the arteries that deliver blood from the heart to other body parts. As expected, coronary artery (*p* =2.63×10 ^-4^) is reported as the top one enriched tissue by the selective expression of associated genes. We also find that the aorta and tibial artery have relatively high enrichment in CAD, probably because CAD associated genes are also selectively expressed among different types of artery. It should be noted that the adipose tissue is prioritized as the third most significant driver tissue, which is consistent with the present studies suggesting the relevance between adipose tissues and CAD [[40], [41], [42]]. Unexpectedly, the nerve is prioritized as the second most significant driver tissue, which we failed to verify in literature survey and requires further research.

#### Rheumatoid arthritis (RA)

RA is a common autoimmune disease mainly attacking the joints. In the selective expression enrichment analysis, two tissues out of the three most significant tissues fall into immune system, spleen and lymphocytes. Unexpectedly, the lung is prioritized as the most significant tissue. However, there have been multiple studies suggesting the lung may play a role in the pathogenesis of RA [43]. There are also studies suggesting lung disease as a major contributor to morbidity and mortality for RA [44]. Some studies suggested lung disease in RA results from targeting of the lung from circulating autoimmunity, but some studies found lung generation of autoimmunity can be present before the onset of joint symptoms^42^. Hence, our results about RA and lung may shed some insights into the relationship between them.

#### Total cholesterol (TC)

For a complex clinical trait, TC, our analysis only detected one significant tissue, liver (*p* = 4.27×10 ^-10^). This is consistent with common knowledge that the liver is responsible for 80% of the endogenous cholesterol synthesis. 30 genes out of 83 TC associated genes are selectively expressed in liver. Among them, several genes have been identified as the causal genes of TC. For example, mutations in PCSK9 has been identified to cause autosomal dominant hypercholesterolemia [45]. Furthermore, recent studies have showed that the monoclonal PCSK9 antibodies could reduce LDL-C in patients with familial and primary hypercholesterolemia alone [[46], [47]].

#### Height

For an anthropometric trait, height, 20 significant tissues are detected, which suggests complex biological mechanisms in the development of human height. However, the most significant tissue is fibroblast (*p* < 10^-16^), which is the most common cells of connective tissue in animals. Consistent with our results, multiple studies, two have also reported connective tissue as the most enriched tissue type for height[47], [48]. Notably, there are also multiple tissues with highly significant p-values (*p* < 10^-7^), such as artery, lung, muscle, adipose, which may provide some new insights into the mechanism of height.

### Comparison of driver tissues detected by our proposed method to another method

We also compared the performance of the proposed framework to a recent method for estimating causal tissues by Ongen et al. [9]. The Figure 2 shows the prioritized tissues of both methods in the above five representative diseases/traits. For bipolar disorder (BD), using our approach, the top ten significant tissues fall into brain areas, most of which have been identified to be relevant with BD [[36], [37], [38], [39]] as mentioned above. However, with Ongen’s approach, only three brain tissues are among the top ten significant tissues. The most significant tissue by Ongen’s approach was prostate, which we failed to verify in literature survey. For coronary artery disease (CAD), coronary artery is correctly prioritized as the most significant tissue by our method, while coronary artery is prioritized as third significant tissue by Ongen’s method. For the autoimmune disease RA, the two immune tissues are prioritized in the top three significant tissues by our proposed method. Unexpectedly, with Ongen’s method, the top five significant tissues even include artery and skin. We do not find studies supporting artery and skin are causal tissues of RA. For total cholesterol, the liver is correctly enriched as most likely causal tissue by both methods. For another complex trait, our results show the fibroblast cells as the most significant causal cells of height, while Ongen’s results suggest the putamen as most significant driver tissues of height. However, in literature few studies investigate the driver tissues of height and it is difficult to evaluate the results of both methods. In general, our proposed framework produced more biologically sensible driver-tissues for complex diseases/traits than Ongen’s method.

### Tissue-selectivity prioritization enhances detection of susceptibility genes in a post GWAS analysis

Finally, we asked how tissue-selectivity prioritization of a disease can be used to enhance detection of susceptibility genes in a post GWAS analysis. In the schizophrenia benchmark dataset, the selective expression ranking leads to ∼30% different significant genes from the default statistical significance ranking in a conditional gene-based test[20]. Among the different significant genes, a rough *in silico* validation in PubMed shows the selective expression ranking results in more genes implicated in schizophrenia by literature than the statistical significance ranking (*n=*16 *vs.* 10, See details in Supplementary Excel Table 1). Here are some individual detailed examples. In a set of physically close genes, the tissue-selective expression ranking and statistical significance ranking lead to different significant genes, DRD2 and MIR4301, respectively. The DRD2 gene is selectively expressed in multiple prioritized pathogenic tissues [including Brain-Anteriorcingulatecortex, Brain-Cortex, Brain-Putamen(basalganglia) and Brain-Spinalcord] and has over 100 papers co-mentioning the gene and schizophrenia in their titles or abstracts in PubMed database. In contrast, there is no paper suggesting MIR4301’s relatedness to schizophrenia. In another set of physically close genes, the tissue-selective expression ranking and statistical significance ranking lead to different significant genes, MIR137HG and MIR2682 respectively. MIR137HG is specifically expressed in above multiple prioritized tissues for schizophrenia [including Brain-Cerebellum, Brain-Anteriorcingulatecortex, and Brain-Hypothalamus]. MIR2682 even had no expression values in any tissues of the GTEx dataset. Moreover, there have been multiple papers (e.g., [49, 50]) suggesting MIR137HG’s contribution to schizophrenia. A bioinformatics prediction by TargetScan (http://www.targetscan.org) suggests this miRNA gene targets multiple empirically validated candidate genes of schizophrenia, e.g., CACNA1C, ZNF804A and TCF4. Among the significant genes based on the statistical significance ranking, ABCB1 has the largest number of PubMed hits. This gene also has selective expression in several prioritized tissues, including Brain-Anteriorcingulatecortex, Brain-FrontalCortex(BA9), and Brain-Substantianigra. Its conditional gene-based association *p*-value according to the tissue-selective expression ranking is suggestively significant *p*=8.7×10^-6^.

In the five representative diseases/traits, the tissue-selective expression ranking consistently results in more previously reported genes than the statistical significance ranking in the conditional gene-based association analysis (See details in Supplementary Excel Table 1). Here are some interesting individual examples. TNF has a significant p-value for rheumatoid arthritis, 3.32×10 ^-25^, according to the tissue-selective expression ranking while it only has a p-value 1.0 according to the statistical significance ranking. The TNF is specifically expressed in immune-related tissues and cells and there are over 100 papers co-mentioning the gene and rheumatoid arthritis in the titles or abstracts in PubMed database. What’s more, TNF-α has been identified as a key molecule in the control of the inflammatory changes that occur in the RA synovium [PMID: 11934972] and the approach of targeting TNF-α has considerably improved the success in the treatment of RA [PMID: 25651945]. For bipolar disorder, MAD1L1 is only detected by the tissue-selectivity prioritization-based method, which has been identified as the candidate gene of bipolar disorder by several studies [[51], [52]]. VEGFA is only detected as the candidate gene of CAD by tissue-selective expression ranking, which has over 100 PubMed hits. Many studies have showed VEGF is dysfunctional in CAD and suggested that VEGF could be used as the biomarker and therapy target for CAD [e.g., [52] [53]]. LIN28B has been identified as an associated gene of height by many studies [e.g., [54]]. While the statistical significance ranking failed to detect it, the tissue-selective expression ranking leads to a very significant p-value (5.46×10 ^-36^). For total cholesterol, SORT1 is only detected by the tissue-selective expression ranking and has been identified as the causal gene of lipid levels [55]. These results suggest the tissue-selectivity prioritization by the proposed robust z-score can enhance detection of true susceptibility genes in a post-GWAS analysis.

## Discussion

To our knowledge, this is the first study to explore how tissue selective expression can be integrated into post-GWAS analysis for identifying potential driver-tissues of complex diseases. Because the associated genes from GWAS are unbiased and potential drivers of diseases, this study provides an unbiased way to precisely explore driver-tissues of complex diseases. Not only can it interpret the mechanism of tissue selective pathogen observed in clinical diagnosis, it can also reveal more comprehensive and primarily relevant tissues than clinical diagnosis. Moreover, the high accessibility of resource data may encourage many explorations of tissue selective pathogen of complex diseases in the future. The expression data and GWAS summary data used in our analysis framework can be downloaded from independent public domains for free and no privacy-sensitive data (e.g., genotypes) are needed. We demonstrated this approach works in six complex diseases or traits of different anatomic systems. First, it systematically confirms many known causal tissues of these diseases. For example, the brains areas are always prioritized for schizophrenia and bipolar disorder. The subtle difference between the prioritized brain tissues may also provide some clues for different clinical manifestation of the two brain diseases. The important immune tissues and cell-types, spleen and lymphocyte cells are significant for rheumatoid arthritis. These biological sensible results in the real examples sufficiently validate the hypothesis that susceptibility genes’ tissue selective expression determines diseases’ tissue selectivity. Second, the comprehensive analysis suggested multiple new candidate driver tissues for further validation, e.g., lung for rheumatoid arthritis and artery for height.

The identified tissues will facilitate transcriptomic and molecular genetic studies of complex diseases. Understanding the path from DNA variation to phenotype variation is a long-held mission of genetics. The recent wave of GWAS consistently implicated that most majority variants confer risk for complex diseases through regulating gene expression[56]. To further study the underlying mechanism, we must know the primitive pathogenic tissues. The experiments in irrelevant tissues will waste time and resources. Although our analyses do not provide direct evidence on how sequence variants change the specific expression in patients to cause a disease, it is unlikely that the significant enrichment of associated genes in many biologically sensible tissues occurred just by chance. One can extrapolate that the tissue selectivity of complex diseases is determined by tissue selective expression of susceptibility genes. Probably due to their selective expression, sequence variation at these genes will impose higher impact on the development of complex diseases. For the detailed mechanism, more molecular biological experiments in the prioritized tissues are needed.

Based on the above extrapolation, we also proposed to use the tissue selective expression to prioritize susceptibility genes in post-GWAS analyses. LD is a tricky problem in GWAS for discriminating true susceptibility genes from indirectly associated genes. Li et al (2018) proposed a powerful statistical framework to isolate possible directly associated genes[20]. However, the original analysis was carried out according to a rank of statistical significance assuming the true susceptibility genes had the more significant p-values. But this is not always true due to expression fluctuations. After ranking genes according to their selective expression in the prioritized tissues for a given disease, we reperformed the conditional gene-based analysis with the new rank. It turned out the selective expression ranks from independent resources led to more significant genes supported by literatures for the six representative complex diseases/traits. These results suggest that integration of selective expression can enhance the power of identifying susceptibility genes. We believe this strategy will also work for many other complex diseases. It will be an effective framework to mine new susceptibility genes in the post-GWAS analyses. Again, the high accessibility of resource data will also encourage similar studies in the post-GWAS era.

A prerequisite of the success in our identification of driver tissues and susceptibility genes is the powerful detection of selectively expressed genes in a tissue. In the comparison of different measures for selective expression, we showed that proposed robust z-score led to the highest significance in the biologically sensible tissues for schizophrenia (Table 2). In contrast, other measures led to either less significant p-values or unlikely tissues. This can be explained by the improved power of the proposed robust z-score in detecting multiple selectively expressed tissues (Table 1). The improved performance is attributed to its technical innovations to conquer problems in real data. The structure of tissue samples in GTEx dataset is complicated with two issues. First, the origin structure of the tissue samples is complex. Some tissues have many similar tissues (e.g. different brains regions) while some have few similar tissues. Second, the size of a tissue sample varies from 5 to 564, which leads to different expression estimation precision. These problems in data are challenging the detection of selective expression. The two issues are not unique for the GTEx dataset; and many other expression datasets have the same problems. The proposed robust z-score was designed on a robust regression of ranked expression and a consideration of expression fluctuation. The robust regression produced robust weights to estimate expression mean and standard error of majority tissues, which circumvents arbitrary partition of the majority and minority tissue. These technical innovations led to improved power in detecting multiple selectively expressed tissues and subsequently disease-related tissues, compared to other selective expression metrics.

The current study also provides abundant data demonstrating that transcript-level selective expression is more powerful for prioritizing driver tissues of complex diseases than that of gene level. Although this looks apparent, few studies have investigated the details. Previous studies often used gene-level selective expression to look into the tissue selective pathogen [2, 14]. In the present study, we show that the usage of transcript level expression (after Bonferroni correction for multiple isoforms) identifies more selectively expressed genes at every tissue [Table 2]. Moreover, the transcript-level selective expression increases the significance p-value for enrichment from 5.8**×**10^-6^ to 5.7**×**10^-14^ at the top tissue of schizophrenia. The comparison suggests studies on transcriptome of complex diseases should pay more attention to transcript level expression. Otherwise, many important expression patterns of susceptibility genes may be overlooked. Moreover, our results also suggest lowly expressed genes or transcripts may be also important for complex diseases when they have large selective expression. The inclusion of lowly expressed genes also led to more significant p-values in the prioritization of driver tissues of schizophrenia (Table 2)

In the five representative examples, we showed the proposed method outperformed the Ongen’s method for estimating causal tissues. The two approaches have different strategies and rationales. Our proposed method is built on the hypothesis that tissue selective expression of susceptibility genes determines the pathogenic selectivity of complex diseases. The powerful gene level association and precise transcript level selective expression are used to estimate the driver tissues. In contrast, the Ongen’s method resorts to an overlap between eQTL in a tissue and GWAS associated variants to estimate the driver tissue. Because of multiple testing burden and small effect size, the variant level association analysis is generally less powerful than the gene level association analysis[57]. Therefore, in theory, the proposed method is more powerful than the Ongen’s. This is also proved in all the tested real examples.

Due to lack of data, the selective expression profiles in 50 tissues are far from complete and the information of developmental stages are lacked either, which is a limitation of the present study. For example, a liver has lobes, surfaces and impressions. In GTEx dataset, liver has no sub-tissues. Due to lack precise tissues, some estimated driver tissues may be still rough in our analysis. Probably due to the same reason, some promising candidate genes have no significant selective expression in any of the prioritized tissues of a disease. The significance in the enrichment analysis may be increased when expression of more related sub-tissues at suitable developmental age is available. However, as more and more expression data are accumulating, this limitation is diminishing. The tissue selective expression will become a powerful resource for identifying driver tissues, developmental stage and new susceptibility genes of human diseases.

## Methods

### The proposed robust measure of tissue selective expression

There are *N* different tissues, and each tissue has multiple transcriptomes. A gene (or transcript) has expression means and standard errors (SE) at the *N* tissues, y_1_,…, y_*N*_ and *s*_1_,…, *s*_*N*_. Assume majority expression values approximately follow a certain distribution (say, normal distribution, or uniform distribution) while a minority of values deviate from the majority due to selective expression.

We notice that the seemingly mussy values in the majority group can often approximately form a line after sorting (See illustration in Supplementary Figure S1). In contrast, the selective expression values will deviate from the line. In addition, as expression means of a gene in tissues with smaller SEs are often more reliable than that with larger SEs, we extended the Huber robust linear regression [58] to weight the expression deviation and reliability. The Huber regression is particularly efficient to outliers in the response variable than other alternative approaches[59, 60], so it can effectively down-weight the expression values deviating from the majority and having large SEs. The regression framework produces smaller weights for the expression values with larger deviation from the fitted line and larger SEs:

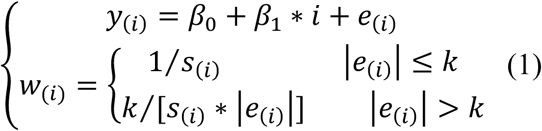

where, *β*_0_ and *β*_1_ are the regression parameters, *i* ∈ [1…*N*] is the rank of a gene expression in the *N* tissues. The y_(*i*)_ denotes the *i*_th_ expression mean in an ascendingly sorted list, and *e*_(*i*)_ denotes the residual. When each tissue only has one subject, *y_i_*, is the expression value of the subject and *s_i_* is set to be 1. The *w*_(*i*)_ is a weight of y_(*i*)_. The *k* is a tuning constant and is equal to 1.345**×** standard deviation of the weighted residuals[58]. The iteratively reweighted least-square procedure of robust linear regression is used to generate the converged weights, *w*_1_,…, *w*_*N*_.

The converged weights are standardized,

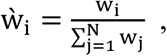

and are used to produce a robust mean,

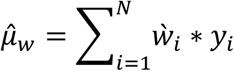

and a robust standard deviation,

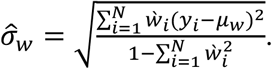

The proposed robust z-score for selective expression at tissue *i* is defined as:

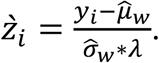

The 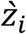 uantifies the expression deviation from the homogenous majority expression values. The λ is a constant factor to adjust the p-values to follow uniform distribution for hypothesis test. Extensive simulations suggested that an empirical factor of 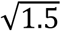 led to approximately uniformly distributed p-values (Figure S2 and S3). The p-value is then approximated based on the standard normal distribution,

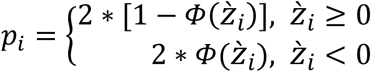, where *Φ*(*x*) is the cumulative distribution function of the standard normal distribution.

### Estimated drive-tissue of diseases by selective expression of disease-associated genes

The driver tissues of a disease are estimated by enrichment analysis of selective expression among genes associated with the disease. The underlying assumption is that tissue selective expression of associated or susceptibility genes of a disease determines the tissue where complex diseases happen primarily, which are called driver tissues. Therefore, a cause tissue is very likely to have an enrichment of selective expression by the susceptibility genes of a disease. We use hypergeometric distribution to evaluate the enrichment. Let’s define that a disease having *m* associated genes. Given a p-value cutoff *c*(=10^-6^), there are *m*_*c*_ genes with significant selective-expression in a tissue among the *m* genes, (*p*_*i*_ ≤ *c*). Let *T*_*c*_ denote the number of genes with selective expression in a tissue one whole genome according to *c* as well (*p*_*i*_ ≤ *c*). *T* denotes the number of all genes having expression values. The enrichment *p* value is calculated by the probability mass function of hypergeometric distribution

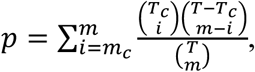

Where 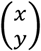 denotes the combinational formula. A significant p-value suggest many associated genes of a disease have selective expression in the tested tissue, indicating the possible driver tissue of the disease.

For a gene with multiple selective expression p-values because of multiple transcripts in a tissue, the smallest p-value of transcripts after Bonferroni correction is used to determine significant selective expression.

### Rank genes by tissue-selective expression in driver tissues

The selective expression in the prioritized tissues is then used to rank candidate genes for a given disease. Assume a disease has *g* significant estimated driver tissues, with the enrichment p-values *p*_1_,…, *p*_*g*_. The ranking score of a gene *j* is

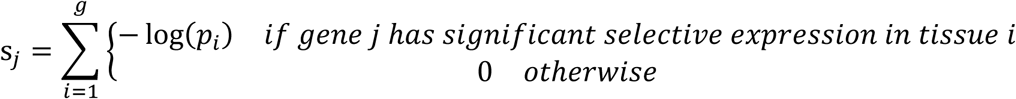

The a gene with selective expression p-value (*p*_*i*_ ≤ *c*) of in a tissue is defined as significant p-value. A gene having selective expression in all prioritized tissues can be given the highest score while a gene having no selective expression in any prioritized tissues is given zero score. The higher score, the more likely to be a susceptibility gene. The ranking score is then used to determine the order in a conditional gene-based association analysis [20]. Genes with higher ranking scores are given higher priority to enter the conditional procedure, which will have a higher chance of getting smaller p-values.

The robust z-score approach, enrichment analysis and gene-ranking score have been implemented into an online tool (http://grass.cgs.hku.hk:8080/dirse/) to facilitate identification of tissue-selective pathogen and susceptibility genes of complex diseases.

### Conditional gene-based based association analysis with GWAS p-values

The conditional gene-based association test, which has an advantage of excluding indirectly associated genes, on KGG (http://grass.cgs.hku.hk/limx/kgg/ [61]) is used to produce disease-associated genes by GWAS summary statistics. For a given disease, the GWAS p-values of sequence variants were input into KGG. The variants within upstream and downstream 5 kb of a gene were assigned onto the gene according to the gene-model, RefGene hg19. Prior to the conditional gene-based association test, an unconditional gene-based test was carried out to detect significant genes by ECS on KGG. The phased genotypes (Phase3 v5 Shapeit2) of EUR panel in 1000 Genomes Project [62] were used as reference genotypes to remove redundant association at sequence variants due to linkage disequilibrium (LD) in the gene-based association test. Among the significant genes detected by the unconditional gene-based association test, some genes may only have indirect association with the disease due to their LD with the true susceptibility genes. The conditional gene-based association analysis was then followed to remove the indirectly associated genes on KGG software [61] according to the same reference genotypes. The conditional gene-based association analysis was performed twice via two different entry orders of genes. At the first time, genes with smaller p-values were given higher priority to enter the iteration for conditional gene-based association analysis. At the second time, genes with higher ranking scores of tissue-selectivity were given higher priority to enter the iteration (See the general pipeline in Figure 1). The basic assumption is that genes selectively expressed in the driver tissues of a complex disease are more likely to be susceptibility genes. If the true susceptibility genes enter first, the conditional gene-based association analysis will more effectively remove indirectly associated genes.

### Gene expression datasets and quality control

The normalized expression datasets at gene level and transcript level were downloaded from GTEx project (V7)[63], GTEx_Analysis_2016-01-15_v7_RNASeQCv1.1.8_gene_tpm.gct.gz and GTEx_Analysis_2016-01-15_v7_RSEMv1.2.22_transcript_tpm.txt.gz. The sample sizes of each tissue were very different, ranging from 5 to 564 (Table S2). There were initially 196,520 transcripts and 56205 genes in 53 tissues. The expression values were measured by transcripts per kilobase million (TPM). As the relative measurement, TPM, has been effective for cross-tissue comparison[64], we did not retransform the expression values by other measurements. A series of quality control procedures were carried out. The mean and standard deviation of expression values of multiple genes in each tissue were produced. In the evaluation according to correlation, three tissues (Whole Blood, Pancreas, and Pituitary) had low Pearson correlation with other tissues (Figure S4 and S5) and were excluded. In the calculation of tissue selective expression, genes or transcripts having ≤0.01 TPM in all tissues were excluded. Genes whose Ensembl IDs had no corresponding official HGNC gene symbols were excluded as well. Finally, 131,292 transcripts and 31,659 genes in 50 tissues were retained for the following analysis.

### Produce genes associated with 5 representative complex diseases or traits

We collected the GWAS meta-analysis p-values at single nucleotide polymorphisms (SNPs) of 5 representative complex diseases or traits developing in different biological systems, bipolar disorder [65](a brain disease), rheumatoid arthritis (RA)[66](an autoimmune disease), coronary artery disease (CAD) [67](a cardiovascular disease), total cholesterol [68] (a metabolic trait), height [69](an anthropometric trait). Table S4 lists the sample sizes and downloading links of the datasets. The p-values of SNPs were combined for gene-based association on KGG (See detailed methods in the above section). The detected associated genes (after multiple testing correction) were used to detect potential driver-tissues and to re-identify additional susceptibility genes (See pipeline in Figure 1).

### Comparison of the proposed framework with another method for estimating driver-tissue

We compared our proposed driver tissue identification method to another causal tissue estimating method proposed by Ongen et al. [9] recently, which was developed under different hypothesis. Ongen’s method estimated the causal tissues according to enrichment of active eQTL in tissues. For fair comparison, we selected the all of 41 overlap GTEx tissues between our study and Ongen et al. [9] for causal tissues estimation of the same diseases. However, because Ongen’s method has no publicly available tools, we directly extracted enrichment values from the Supplementary Table S5 of their published paper [9]. According to Ongen et al., the tissues with the enrichment value over the null greater than 1 was consider as the causal tissues for the diseases/traits.

### In silico validation by PubMed search

We used PubMed search function to validate the detected genes for a complex disease. The underlying assumption is that multiple papers co-mentioning a gene and a disease name in the title or abstract may implicate the contribution of the gene to the disease. The more hit papers, the more likelihood the gene is related to the disease. Although this may be crude for a specific gene, it can produce a reliable systematic evaluation when there are many genes. We employed the web application programming interfaces (APIs) of PubMed to execute the search. The search link was, http://eutils.ncbi.nlm.nih.gov/entrez/eutils/esearch.fcgi?db=pubmed&term=“DiseaseNames(inlcuding homonymies)”[tiab]%29+AND+“GeneSymbol (including RefSeq mRNA IDs)” [tiab]. The search results included PubMed ID and relevant data of the papers, if available, in extensible markup language (XML).

## Supporting information

## Acknowledgements

This work was funded by National Natural Science Foundation of China (31771401), National Key R&D Program of China(2018YFC0910500). Hong Kong General Research Fund 17124017, 17121414 and TRS T12C-714/14-R. We thank The GTEx Consortium and Genome Aggregation Database to provide access for expression and sequence variants data on thousands of samples from various tissues.

## Conflict of interest

The authors declare that they have no competing interests.

## Author Contributions

Lin Jiang: analyze the data, draft the article and interpretation of data, and develop the methods and tool;

Chao Xue: analyze of the data, draft the article and interpretation of data, and develop the methods and tool;

Shangzhen Chen: analyze of data and design the website; Shen Dai: develop the methods and tool;

Peikai Chen: process and analyze of data;

Pak Chung Sham: contribute to conception and design and revise the manuscript

Haijun Wang: substantially contribute to conception and design, revise it critically for important intellectual content and final approval of the version to be published;

Miaoxin Li: draft the article, develop the methods and tool, substantially contribute to conception and design, revise it critically for important intellectual content and final approval of the version to be published.

## Availability of data and materials

DIRSE, http://grass.cgs.hku.hk:8080/dirse/

KGG, http://grass.cgs.hku.hk/limx/kgg/

KGGSeq, http://grass.cgs.hku.hk/limx/kggseq/

Clinical Genomic Database https://research.nhgri.nih.gov/CGD/download/xls/CGD.xls.gz

COSMIC database, http://cancer.sanger.ac.uk/cosmic/

Genome Aggregation Database, http://gnomad.broadinstitute.org/

GTEx, https://www.gtexportal.org/home/

## References

1. Barshir R, Shwartz O, Smoly IY, Yeger-Lotem E: Comparative analysis of human tissue interactomes reveals factors leading to tissue-specific manifestation of hereditary diseases. PLoS Comput Biol 2014, 10: e1003632.

2. Lage K, Hansen NT, Karlberg EO, Eklund AC, Roque FS, Donahoe PK, Szallasi Z, Jensen TS, Brunak S: A large-scale analysis of tissue-specific pathology and gene expression of human disease genes and complexes. Proc Natl Acad Sci U S A 2008, 105: 20870–20875.

3. Greene CS, Krishnan A, Wong AK, Ricciotti E, Zelaya RA, Himmelstein DS, Zhang R, Hartmann BM, Zaslavsky E, Sealfon SC, et al: Understanding multicellular function and disease with human tissue-specific networks. Nat Genet 2015, 47: 569–576.

4. Schaid DJ, Chen W, Larson NB: From genome-wide associations to candidate causal variants by statistical fine-mapping. Nat Rev Genet 2018, 19: 491–504.

5. DeLisi LE, Szulc KU, Bertisch HC, Majcher M, Brown K: Understanding structural brain changes in schizophrenia. Dialogues Clin Neurosci 2006, 8: 71–78.

6. Calderon D, Bhaskar A, Knowles DA, Golan D, Raj T, Fu AQ, Pritchard JK: Inferring Relevant Cell Types for Complex Traits by Using Single-Cell Gene Expression. Am J Hum Genet 2017, 101: 686–699.

7. Consortium GT, Laboratory DA, Coordinating Center-Analysis Working G, Statistical Methods groups-Analysis Working G, Enhancing Gg, Fund NIHC, Nih/Nci, Nih/Nhgri, Nih/Nimh, Nih/Nida, et al: Genetic effects on gene expression across human tissues. Nature 2017, 550: 204–213.

8. Antanaviciute A, Daly C, Crinnion LA, Markham AF, Watson CM, Bonthron DT, Carr IM: GeneTIER: prioritization of candidate disease genes using tissue-specific gene expression profiles. Bioinformatics 2015, 31: 2728–2735.

9. Ongen H, Brown AA, Delaneau O, Panousis NI, Nica AC, Consortium GT, Dermitzakis ET: Estimating the causal tissues for complex traits and diseases. Nat Genet 2017, 49: 1676–1683.

10. Liang S, Li Y, Be X, Howes S, Liu W: Detecting and profiling tissue-selective genes. Physiol Genomics 2006, 26: 158–162.

11. Kryuchkova-Mostacci N, Robinson-Rechavi M: A benchmark of gene expression tissue-specificity metrics. Brief Bioinform 2017, 18: 205–214.

12. Kadota K, Nishimura S, Bono H, Nakamura S, Hayashizaki Y, Okazaki Y, Takahashi K: Detection of genes with tissue-specific expression patterns using Akaike’s information criterion procedure. In Physiol Genomics, vol. 12. pp. 251–259; 2003:251-259.

13. Kadota K, Ye J, Nakai Y, Terada T, Shimizu K: ROKU: a novel method for identification of tissue-specific genes. BMC Bioinformatics 2006, 7: 294.

14. Xiao SJ, Zhang C, Zou Q, Ji ZL: TiSGeD: a database for tissue-specific genes. Bioinformatics 2010, 26: 1273–1275.

15. Mele M, Ferreira PG, Reverter F, DeLuca DS, Monlong J, Sammeth M, Young TR, Goldmann JM, Pervouchine DD, Sullivan TJ, et al: Human genomics. The human transcriptome across tissues and individuals. Science 2015, 348: 660–665.

16. Ogasawara O, Otsuji M, Watanabe K, Iizuka T, Tamura T, Hishiki T, Kawamoto S, Okubo K: BodyMap-Xs: anatomical breakdown of 17 million animal ESTs for cross-species comparison of gene expression. Nucleic Acids Res 2006, 34: D628–631.

17. Liu X, Yu X, Zack DJ, Zhu H, Qian J: TiGER: a database for tissue-specific gene expression and regulation. BMC Bioinformatics 2008, 9: 271.

18. Li X, Kim Y, Tsang EK, Davis JR, Damani FN, Chiang C, Hess GT, Zappala Z, Strober BJ, Scott AJ, et al: The impact of rare variation on gene expression across tissues. Nature 2017, 550: 239–243.

19. Tan MH, Li Q, Shanmugam R, Piskol R, Kohler J, Young AN, Liu KI, Zhang R, Ramaswami G, Ariyoshi K, et al: Dynamic landscape and regulation of RNA editing in mammals. Nature 2017, 550: 249–254.

20. Li M, Jiang L, Mak TSH, Kwan JSH, Xue C, Chen P, Leung HC, Cui L, Li T, Sham PC: A powerful conditional gene-based association approach implicated functionally important genes for schizophrenia. Bioinformatics 2018.

21. Sheikh F, Lyon RC, Chen J: Functions of myosin light chain-2 (MYL2) in cardiac muscle and disease. Gene 2015, 569: 14–20.

22. Yuan CC, Kazmierczak K, Liang J, Zhou Z, Yadav S, Gomes AV, Irving TC, Szczesna-Cordary D: Sarcomeric perturbations of myosin motors lead to dilated cardiomyopathy in genetically modified MYL2 mice. Proc Natl Acad Sci U S A 2018, 115: E2338–E2347.

23. Peng W, Li M, Li H, Tang K, Zhuang J, Zhang J, Xiao J, Jiang H, Li D, Yu Y, et al: Dysfunction of Myosin Light-Chain 4 (MYL4) Leads to Heritable Atrial Cardiomyopathy With Electrical, Contractile, and Structural Components: Evidence From Genetically-Engineered Rats. J Am Heart Assoc 2017, 6.

24. Amiel J, Rio M, de Pontual L, Redon R, Malan V, Boddaert N, Plouin P, Carter NP, Lyonnet S, Munnich A, Colleaux L: Mutations in TCF4, encoding a class I basic helix-loop-helix transcription factor, are responsible for Pitt-Hopkins syndrome, a severe epileptic encephalopathy associated with autonomic dysfunction. Am J Hum Genet 2007, 80: 988–993.

25. Hamdan FF, Srour M, Capo-Chichi JM, Daoud H, Nassif C, Patry L, Massicotte C, Ambalavanan A, Spiegelman D, Diallo O, et al: De novo mutations in moderate or severe intellectual disability. PLoS Genet 2014, 10: e1004772.

26. Basmanav FB, Forstner AJ, Fier H, Herms S, Meier S, Degenhardt F, Hoffmann P, Barth S, Fricker N, Strohmaier J, et al: Investigation of the role of TCF4 rare sequence variants in schizophrenia. Am J Med Genet B Neuropsychiatr Genet 2015, 168B:354–362.

27. Schizophrenia Working Group of the Psychiatric Genomics C: Biological insights from 108 schizophrenia-associated genetic loci. Nature 2014, 511: 421–427.

28. Fornito A, Yucel M, Dean B, Wood SJ, Pantelis C: Anatomical abnormalities of the anterior cingulate cortex in schizophrenia: bridging the gap between neuroimaging and neuropathology. Schizophr Bull 2009, 35: 973–993.

29. Andreasen NC, Pierson R: The role of the cerebellum in schizophrenia. Biol Psychiatry 2008, 64: 81–88.

30. Knable MB, Weinberger DR: Dopamine, the prefrontal cortex and schizophrenia. J Psychopharmacol 1997, 11: 123–131.

31. Bjorkquist OA, Olsen EK, Nelson BD, Herbener ES: Altered amygdala-prefrontal connectivity during emotion perception in schizophrenia. Schizophr Res 2016, 175: 35–41.

32. Allen KM, Fung SJ, Weickert CS: Cell proliferation is reduced in the hippocampus in schizophrenia. Aust N Z J Psychiatry 2016, 50: 473–480.

33. Moberget T, Doan NT, Alnaes D, Kaufmann T, Cordova-Palomera A, Lagerberg TV, Diedrichsen J, Schwarz E, Zink M, Eisenacher S, et al: Cerebellar volume and cerebellocerebral structural covariance in schizophrenia: a multisite mega-analysis of 983 patients and 1349 healthy controls. Mol Psychiatry 2018, 23: 1512–1520.

34. Quednow BB, Brzozka MM, Rossner MJ: Transcription factor 4 (TCF4) and schizophrenia: integrating the animal and the human perspective. Cell Mol Life Sci 2014, 71: 2815–2835.

35. He K, An Z, Wang Q, Li T, Li Z, Chen J, Li W, Wang T, Ji J, Feng G, et al: CACNA1C, schizophrenia and major depressive disorder in the Han Chinese population. Br J Psychiatry 2014, 204: 36–39.

36. Phillips JR, Hewedi DH, Eissa AM, Moustafa AA: The cerebellum and psychiatric disorders. Front Public Health 2015, 3: 66.

37. Frey BN, Andreazza AC, Nery FG, Martins MR, Quevedo J, Soares JC, Kapczinski F: The role of hippocampus in the pathophysiology of bipolar disorder. Behav Pharmacol 2007, 18: 419–430.

38. Daban C, Vieta E, Mackin P, Young AH: Hypothalamic-pituitary-adrenal axis and bipolar disorder. Psychiatr Clin North Am 2005, 28: 469–480.

39. Hwang J, Lyoo IK, Dager SR, Friedman SD, Oh JS, Lee JY, Kim SJ, Dunner DL, Renshaw PF: Basal ganglia shape alterations in bipolar disorder. Am J Psychiatry 2006, 163: 276–285.

40. Guauque-Olarte S, Gaudreault N, Piche ME, Fournier D, Mauriege P, Mathieu P, Bosse Y: The transcriptome of human epicardial, mediastinal and subcutaneous adipose tissues in men with coronary artery disease. PLoS One 2011, 6: e19908.

41. Ahn SG, Lim HS, Joe DY, Kang SJ, Choi BJ, Choi SY, Yoon MH, Hwang GS, Tahk SJ, Shin JH: Relationship of epicardial adipose tissue by echocardiography to coronary artery disease. Heart 2008, 94: e7.

42. Golia E, Limongelli G, Natale F, Fimiani F, Maddaloni V, Russo PE, Riegler L, Bianchi R, Crisci M, Palma GD, et al: Adipose tissue and vascular inflammation in coronary artery disease. World J Cardiol 2014, 6: 539–554.

43. Demoruelle MK, Solomon JJ, Fischer A, Deane KD: The lung may play a role in the pathogenesis of rheumatoid arthritis. Int J Clin Rheumtol 2014, 9: 295–309.

44. Kim EJ, Collard HR, King TE, Jr.: Rheumatoid arthritis-associated interstitial lung disease: the relevance of histopathologic and radiographic pattern. Chest 2009, 136: 1397–1405.

45. Abifadel M, Varret M, Rabes JP, Allard D, Ouguerram K, Devillers M, Cruaud C, Benjannet S, Wickham L, Erlich D, et al: Mutations in PCSK9 cause autosomal dominant hypercholesterolemia. Nat Genet 2003, 34: 154–156.

46. Dias CS, Shaywitz AJ, Wasserman SM, Smith BP, Gao B, Stolman DS, Crispino CP, Smirnakis KV, Emery MG, Colbert A, et al: Effects of AMG 145 on low-density lipoprotein cholesterol levels: results from 2 randomized, double-blind, placebo-controlled, ascending-dose phase 1 studies in healthy volunteers and hypercholesterolemic subjects on statins. J Am Coll Cardiol 2012, 60: 1888–1898.

47. Stein EA, Gipe D, Bergeron J, Gaudet D, Weiss R, Dufour R, Wu R, Pordy R: Effect of a monoclonal antibody to PCSK9, REGN727/SAR236553, to reduce low-density lipoprotein cholesterol in patients with heterozygous familial hypercholesterolaemia on stable statin dose with or without ezetimibe therapy: a phase 2 randomised controlled trial. Lancet 2012, 380: 29–36.

48. Hao X, Zeng P, Zhang S, Zhou X: Identifying and exploiting trait-relevant tissues with multiple functional annotations in genome-wide association studies. PLoS Genet 2018, 14: e1007186.

49. He E, Lozano MAG, Stringer S, Watanabe K, Sakamoto K, den Oudsten F, Koopmans F, Giamberardino SN, Hammerschlag A, Cornelisse LN, et al: MIR137 schizophrenia-associated locus controls synaptic function by regulating synaptogenesis, synapse maturation and synaptic transmission. Hum Mol Genet 2018, 27: 1879–1891.

50. Wright C, Gupta CN, Chen J, Patel V, Calhoun VD, Ehrlich S, Wang L, Bustillo JR, Perrone-Bizzozero NI, Turner JA: Polymorphisms in MIR137HG and microRNA-137-regulated genes influence gray matter structure in schizophrenia. Transl Psychiatry 2016, 6: e724.

51. Ikeda M, Takahashi A, Kamatani Y, Okahisa Y, Kunugi H, Mori N, Sasaki T, Ohmori T, Okamoto Y, Kawasaki H, et al: A genome-wide association study identifies two novel susceptibility loci and trans population polygenicity associated with bipolar disorder. Mol Psychiatry 2018, 23: 639–647.

52. Trost S, Diekhof EK, Mohr H, Vieker H, Kramer B, Wolf C, Keil M, Dechent P, Binder EB, Gruber O: Investigating the Impact of a Genome-Wide Supported Bipolar Risk Variant of MAD1L1 on the Human Reward System. Neuropsychopharmacology 2016, 41: 2679–2687.

53. Marks ECA, Wilkinson TM, Frampton CM, Skelton L, Pilbrow AP, Yandle TG, Pemberton CJ, Doughty RN, Whalley GA, Ellis CJ, et al: Plasma levels of soluble VEGF receptor isoforms, circulating pterins and VEGF system SNPs as prognostic biomarkers in patients with acute coronary syndromes. BMC Cardiovasc Disord 2018, 18: 169.

54. Cousminer DL, Berry DJ, Timpson NJ, Ang W, Thiering E, Byrne EM, Taal HR, Huikari V, Bradfield JP, Kerkhof M, et al: Genome-wide association and longitudinal analyses reveal genetic loci linking pubertal height growth, pubertal timing and childhood adiposity. Hum Mol Genet 2013, 22: 2735–2747.

55. Musunuru K, Strong A, Frank-Kamenetsky M, Lee NE, Ahfeldt T, Sachs KV, Li X, Li H, Kuperwasser N, Ruda VM, et al: From noncoding variant to phenotype via SORT1 at the 1p13 cholesterol locus. Nature 2010, 466: 714–719.

56. Gao L, Uzun Y, Gao P, He B, Ma X, Wang J, Han S, Tan K: Identifying noncoding risk variants using disease-relevant gene regulatory networks. Nat Commun 2018, 9: 702.

57. Li MX, Gui HS, Kwan JS, Sham PC: GATES: a rapid and powerful gene-based association test using extended Simes procedure. Am J Hum Genet 2011, 88: 283–293.

58. Huber PJ, Ronchetti EM: Robust Statistics. 2 edn: John Wiley & Sons Inc.; 2009.

59. Reddy BR, Narayan KL, Pattabhiramacharyulu NC: On Global Stability of Two Mutually Interacting Species with Limited Resources for both the Species Int J Contemp Math Sciences 2011, 6: 401–407.

60. AlmaÖ G: Comparison of Robust Regression Methods in Linear Regression. Int J Contemp Math Sciences 2011, 6: 409–421.

61. Li MX, Sham PC, Cherny SS, Song YQ: A knowledge-based weighting framework to boost the power of genome-wide association studies. PLoS One 2010, 5: e14480.

62. Genomes Project C, Auton A, Brooks LD, Durbin RM, Garrison EP, Kang HM, Korbel JO, Marchini JL, McCarthy S, McVean GA, Abecasis GR: A global reference for human genetic variation. Nature 2015, 526: 68–74.

63. e GP: Enhancing GTEx by bridging the gaps between genotype, gene expression, and disease. Nat Genet 2017, 49: 1664–1670.

64. Li B, Ruotti V, Stewart RM, Thomson JA, Dewey CN: RNA-Seq gene expression estimation with read mapping uncertainty. Bioinformatics 2010, 26: 493–500.

65. Bipolar D, Schizophrenia Working Group of the Psychiatric Genomics Consortium. Electronic address drve, Bipolar D, Schizophrenia Working Group of the Psychiatric Genomics C: Genomic Dissection of Bipolar Disorder and Schizophrenia, Including 28 Subphenotypes. Cell 2018, 173: 1705–1715 e1716.

66. Okada Y, Wu D, Trynka G, Raj T, Terao C, Ikari K, Kochi Y, Ohmura K, Suzuki A, Yoshida S, et al: Genetics of rheumatoid arthritis contributes to biology and drug discovery. Nature 2014, 506: 376–381.

67. Nikpay M, Goel A, Won HH, Hall LM, Willenborg C, Kanoni S, Saleheen D, Kyriakou T, Nelson CP, Hopewell JC, et al: A comprehensive 1,000 Genomes-based genome-wide association meta-analysis of coronary artery disease. Nat Genet 2015, 47: 1121–1130.

68. Willer CJ, Schmidt EM, Sengupta S, Peloso GM, Gustafsson S, Kanoni S, Ganna A, Chen J, Buchkovich ML, Mora S, et al: Discovery and refinement of loci associated with lipid levels. Nat Genet 2013, 45: 1274–1283.

69. Wood AR, Esko T, Yang J, Vedantam S, Pers TH, Gustafsson S, Chu AY, Estrada K, Luan J, Kutalik Z, et al: Defining the role of common variation in the genomic and biological architecture of adult human height. Nat Genet 2014, 46: 1173–1186.

