## Supplementary material for "Estimating driver-tissues by robust selective expression of genes associated with complex diseases or traits"

**Supplementary Text:**

1. **Calculate weighted mean and standard deviation**

The iterative approach finally produces positive weights for each expression value, $w_{1},\ldots, w_{N}$. The larger deviation from the fitted line, the smaller weights will be given at the expression values. The weights are then standardized to be reliability weights:

$\grave{w}_{i}=\frac{w_{i}}{\sum_{j=1}^{N} w_{j}}$.

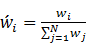

And the reliability weights are then used to calculate weighted mean ($\mu_{w}$) and weighted unbiased standard deviation ($\sigma_{w}$) of the expression values:

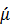

$$\mu_{w}=\sum_{i=1}^{N} \grave{w}_{i}*y_{i}$$

$\sigma_{w}=\sqrt{\frac{\sum_{i=1}^{N} \grave{w}_{i}(y_{i}-\mu_{w})^{2}}{1-\sum_{i=1}^{N} \grave{w}_{i}^{2}}}$.

2. **Simulation studies to evaluate type 1 error and power of the robust z-score approach**

We performed computer simulation to evaluate the type 1 error and the power of the robust z-score approach. For evaluation of type 1 error, *N* expression values under normal distribution were produced for *N* tissues. This way simulated null hypothesis in which no tissues had specifically high or low expression among the *N* tissues. The robust z-score approach was used to produce tissue specific expression scores and to calculate corresponding p-values. Under the null hypothesis, the tissue specific expression scores approximately follow standard normal distribution and the p-values were expected to follow uniform distribution approximately. We repeated the simulation 100,000 to obtain *N*∙100,000 *p* values. The ${-log}_{10} p$ was used to draw a quantile-quantile(QQ) plot under uniform distribution. For power evaluation, *k* ($\ll N$) tissues were assumed to have expression with *d*-fold standard deviation deviating from the mean. Again, the simulation was repeated 100,000 times and the *p*-values for tissue-specific expression were calculated by the proposed approach. Given a p-value cutoff for Bonferroni correction at a family-wise error rate 0.05, the power of a test was defined as the proportion of simulations with significant p-values in the specifically expressed tissues out of the 100,000 simulations.

3. **Implementation**

We have implemented and released the proposed methods into an user-friendly online tool for estimating driver tissues and curated tissue specific expression data at DIRSE website, http://grass.cgs.hku.hk:8080/dirse/. The tool is encoded by Java Server Pages, running on a web server. Users can input a set of disease associated genes and choose the interested tissues or cell-types. The tool will calculate selective expression of genes at every tissue and estimate driver tissues by the selective expression of input genes. Two parameters (pCut and TPMCut) on the tool can be set to define selectively expressed genes and exclude lowly expressed genes and transcripts. In addition, users can input either a single gene through a web browser to query specific expression at transcripts in 50 tissues. See details of the tool on the webpage, http://grass.cgs.hku.hk:8080/dirse/tutorial/.

**Supplementary Figures**

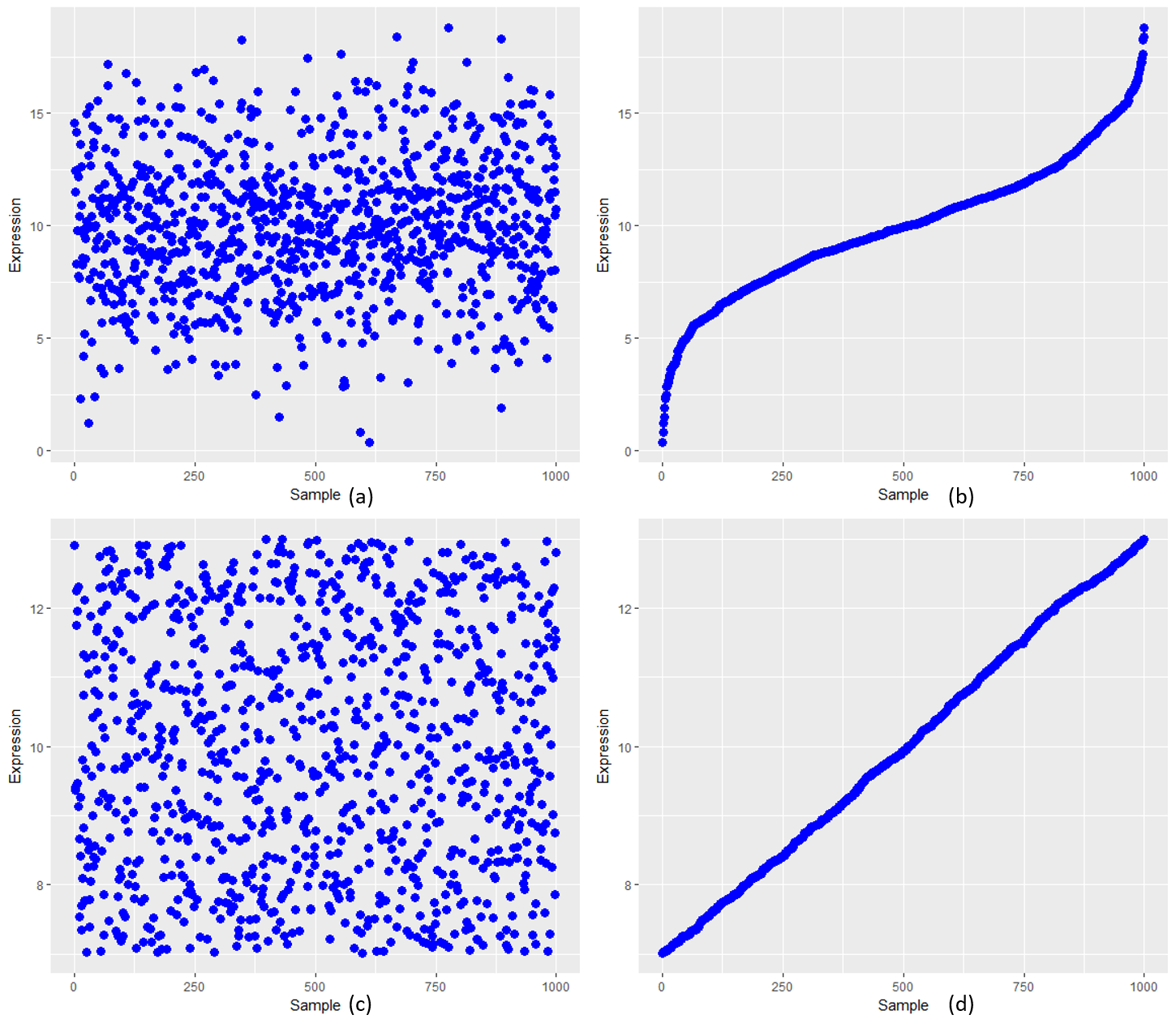

Figure S1: Illustration of approximate linear features of normalized expression values after sorting. a) the expression values prior to sorting under normal distribution b) the expression values after sorting under normal distribution

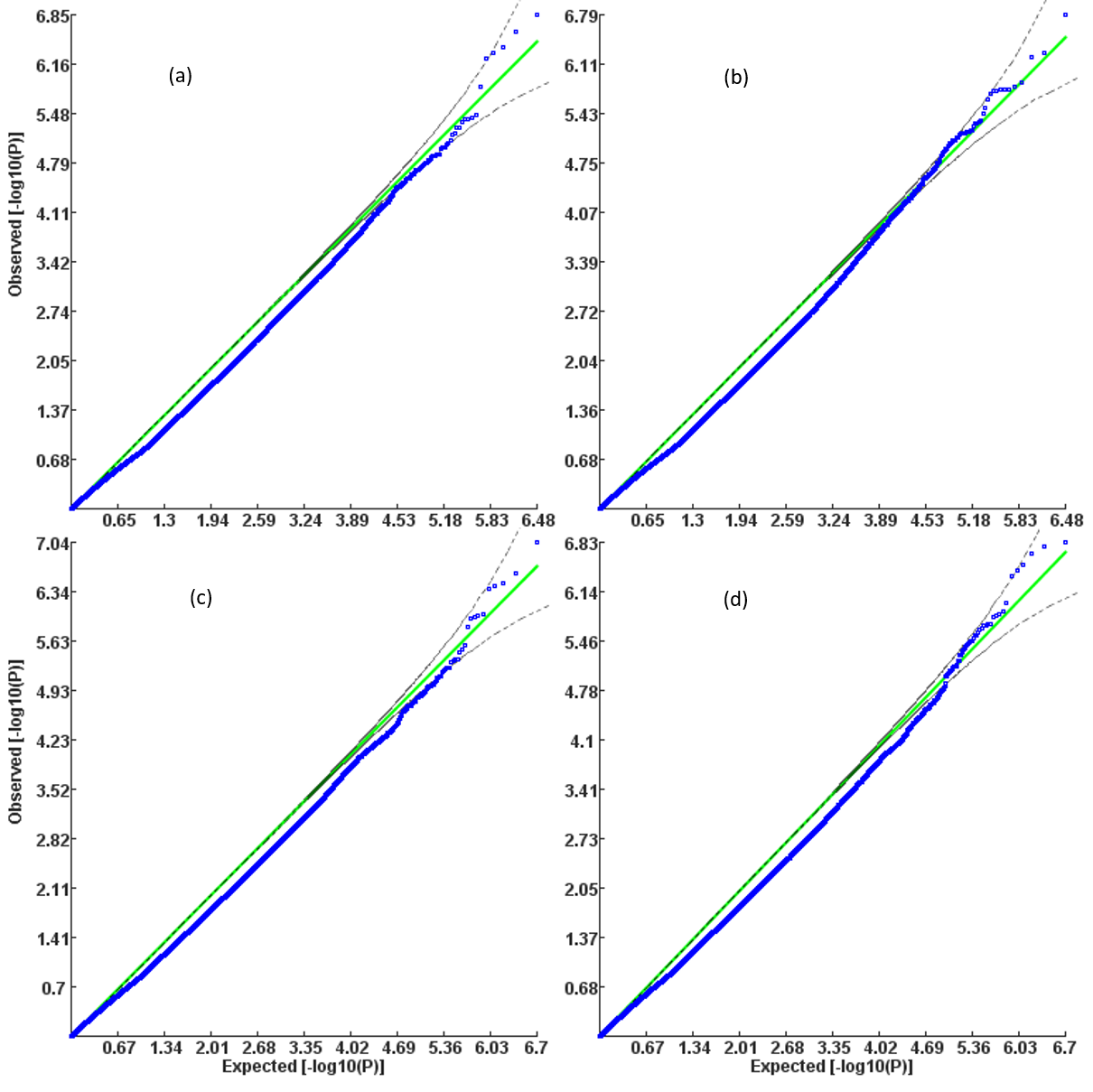

Figure S2: QQ plot of for detecting tissue-specific expression under simulated normally distributed expression values a)the distribution of expression: N(1,1), number of tissues:30, and sample size of each tissue [70,100]; b)the distribution of expression: N(10,5), number of tissues:30, and sample size of each tissue [70,100]; c)the distribution of expression: N(10,10), number of tissues:50, and sample size of each tissue [70,100], sample size of proposed robust z-score; d)the distribution of expression: N(10,10), number of tissues:50, and sample size of each tissue [300,400]. In each scenario, 100,000 datasets were simulated for the analysis.

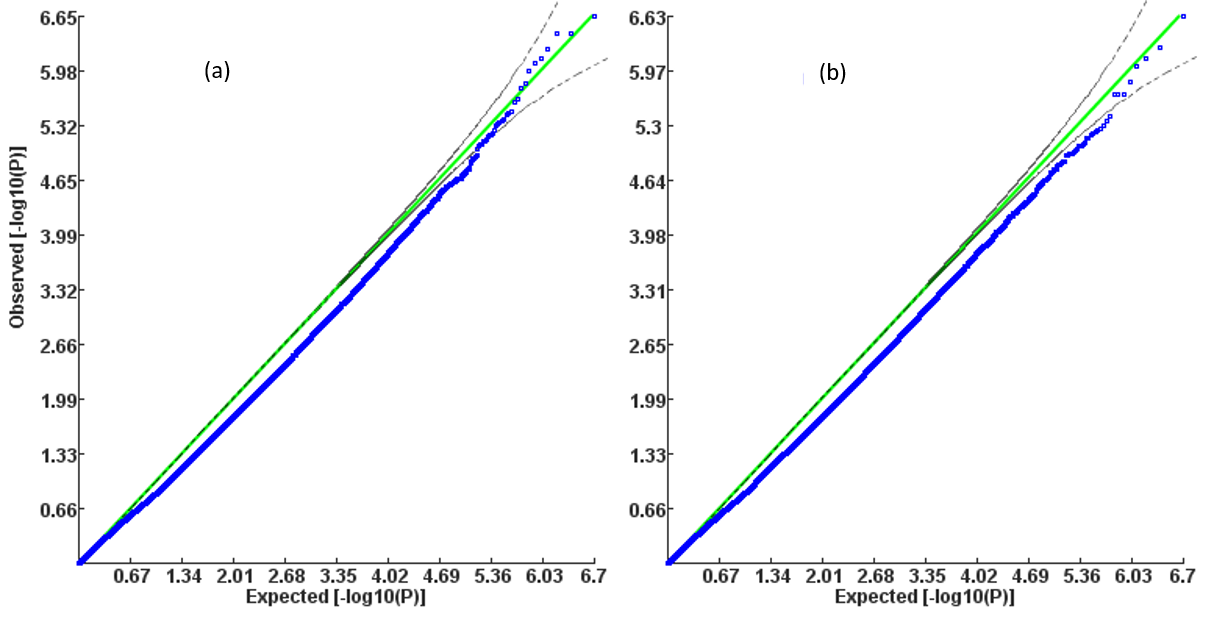

Figure S3: QQ plot of for detecting tissue-specific expression under simulated uniformly distributed expression values a)the distribution of expression: U(5,15), number of tissues:50, and sample size of each tissue [70,100]; b)the distribution of expression: U(25,75), number of tissues:50, and sample size of each tissue [300,400]. In each scenario, 100,000 datasets were simulated for the analysis.

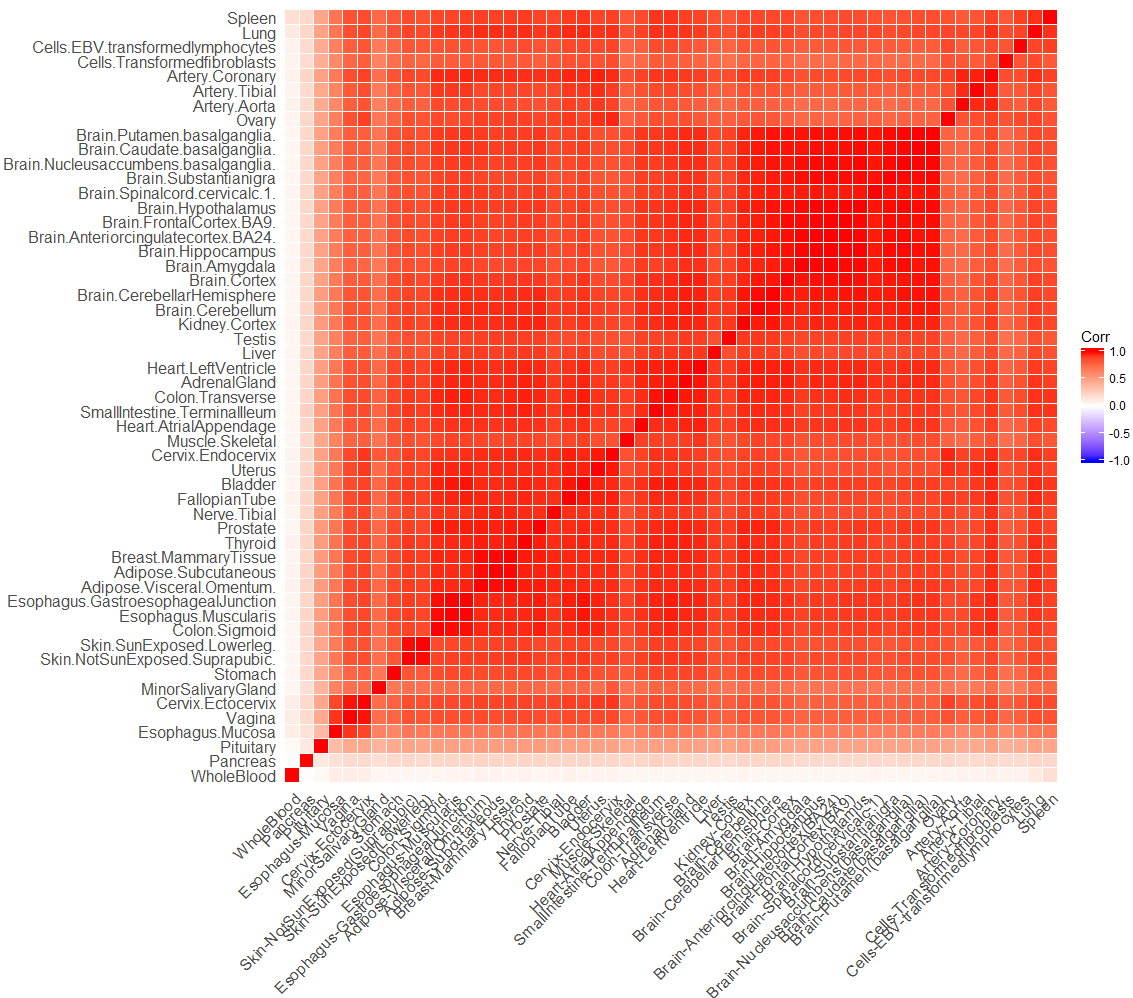

Figure S4: Pearson correlation of the tissues according to the TPM scores at genes.

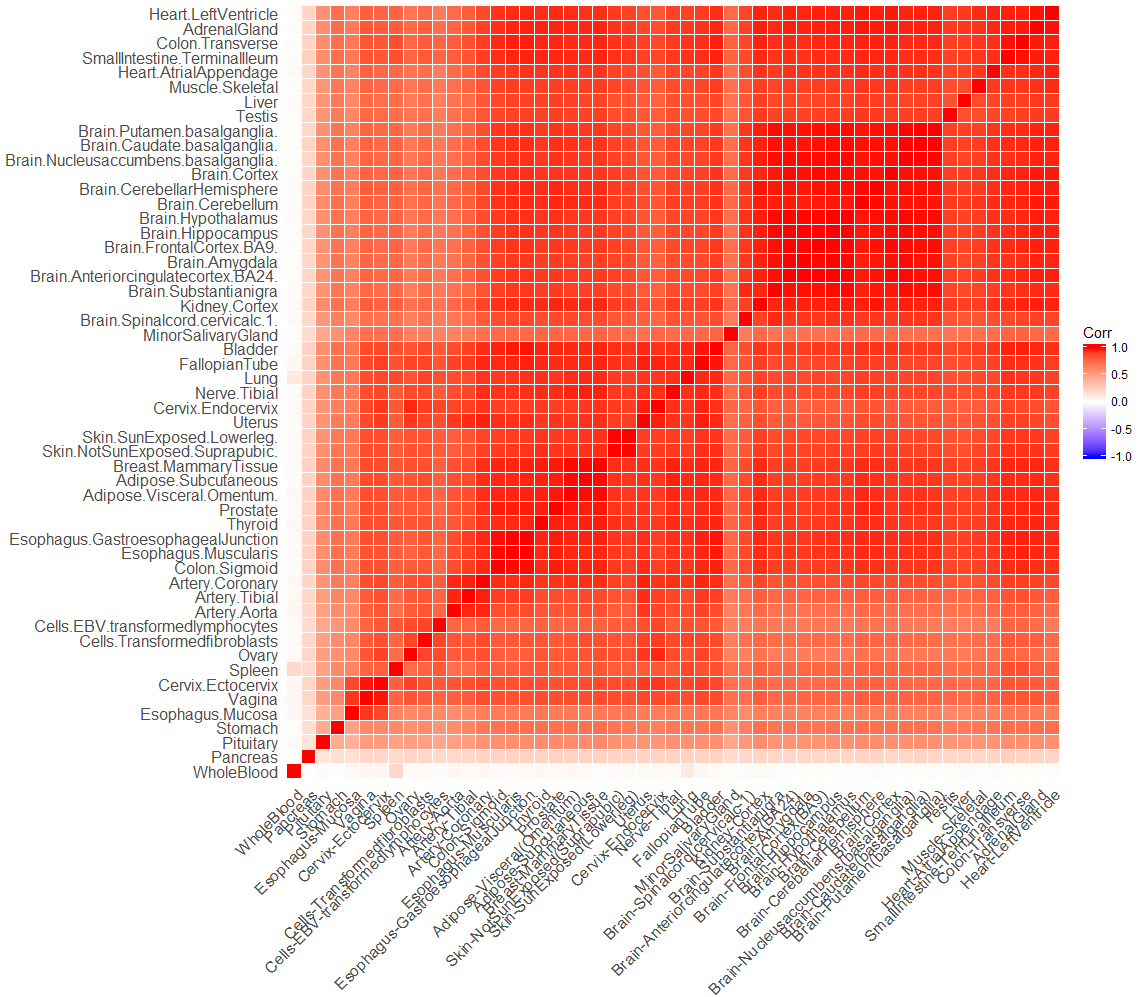

Figure S5: Pearson correlation of the tissues according to the TPM scores at transcripts.

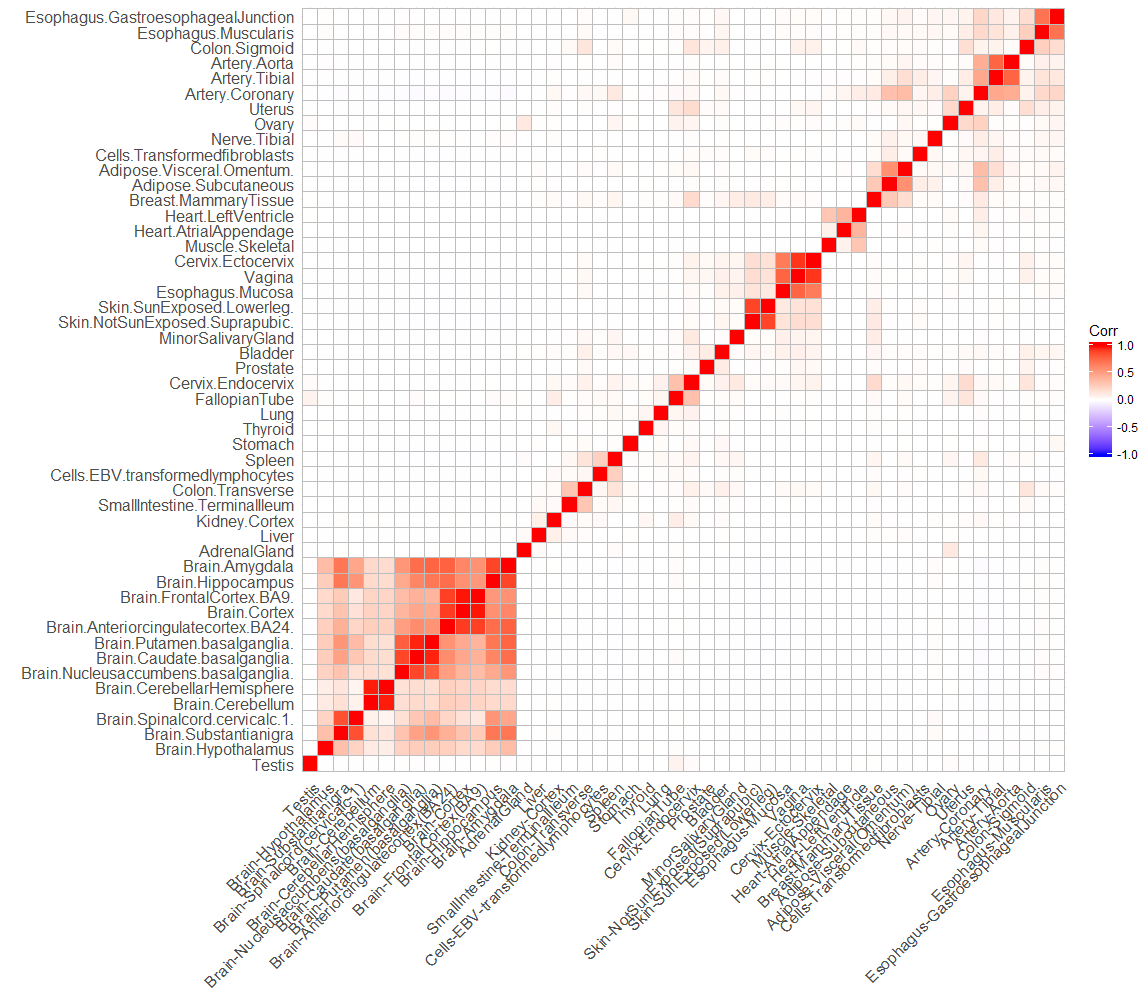

Figure S6: Pearson correlation of the tissues calculated by the robust-z scores at genes.

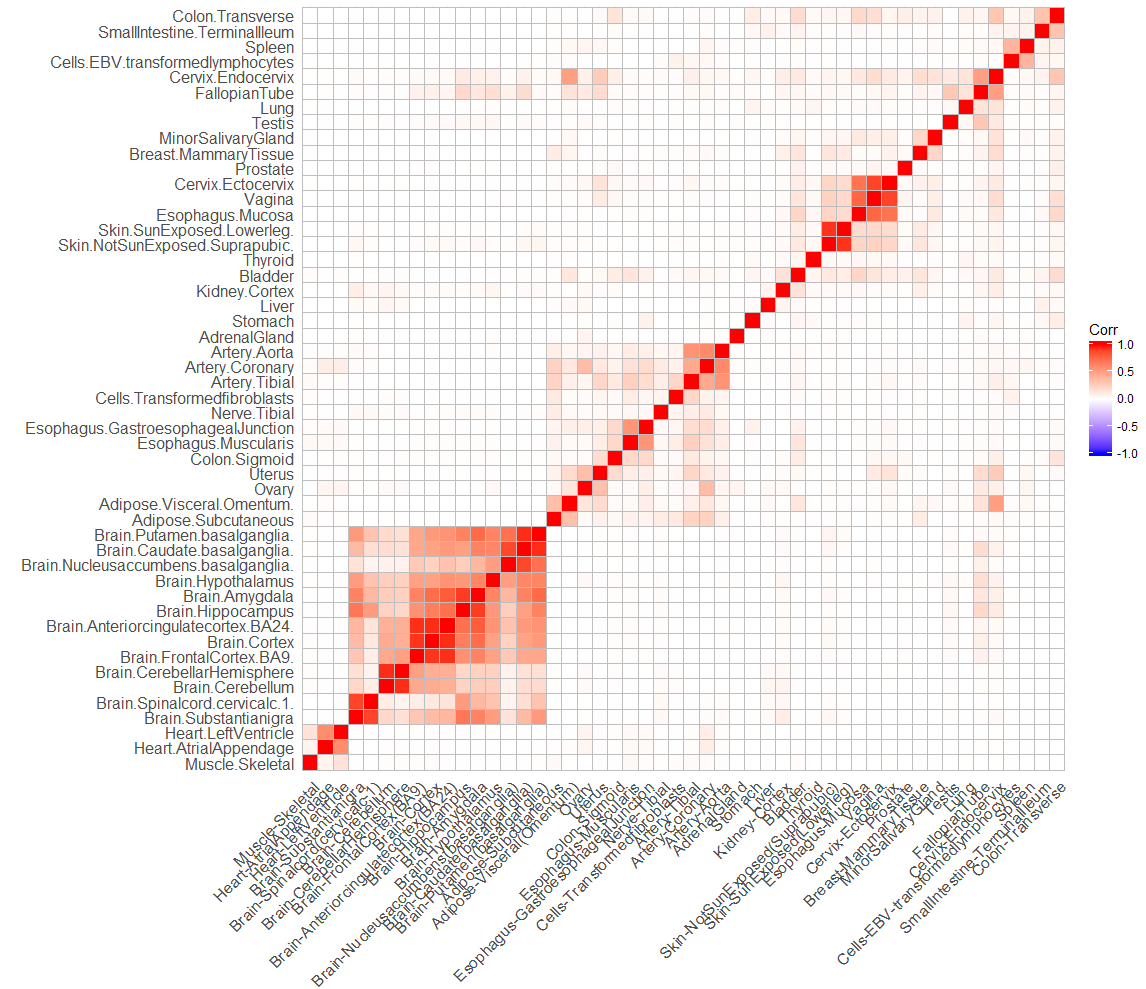

Figure S7: Pearson correlation of the tissues calculated by the robust-z scores at transcripts.

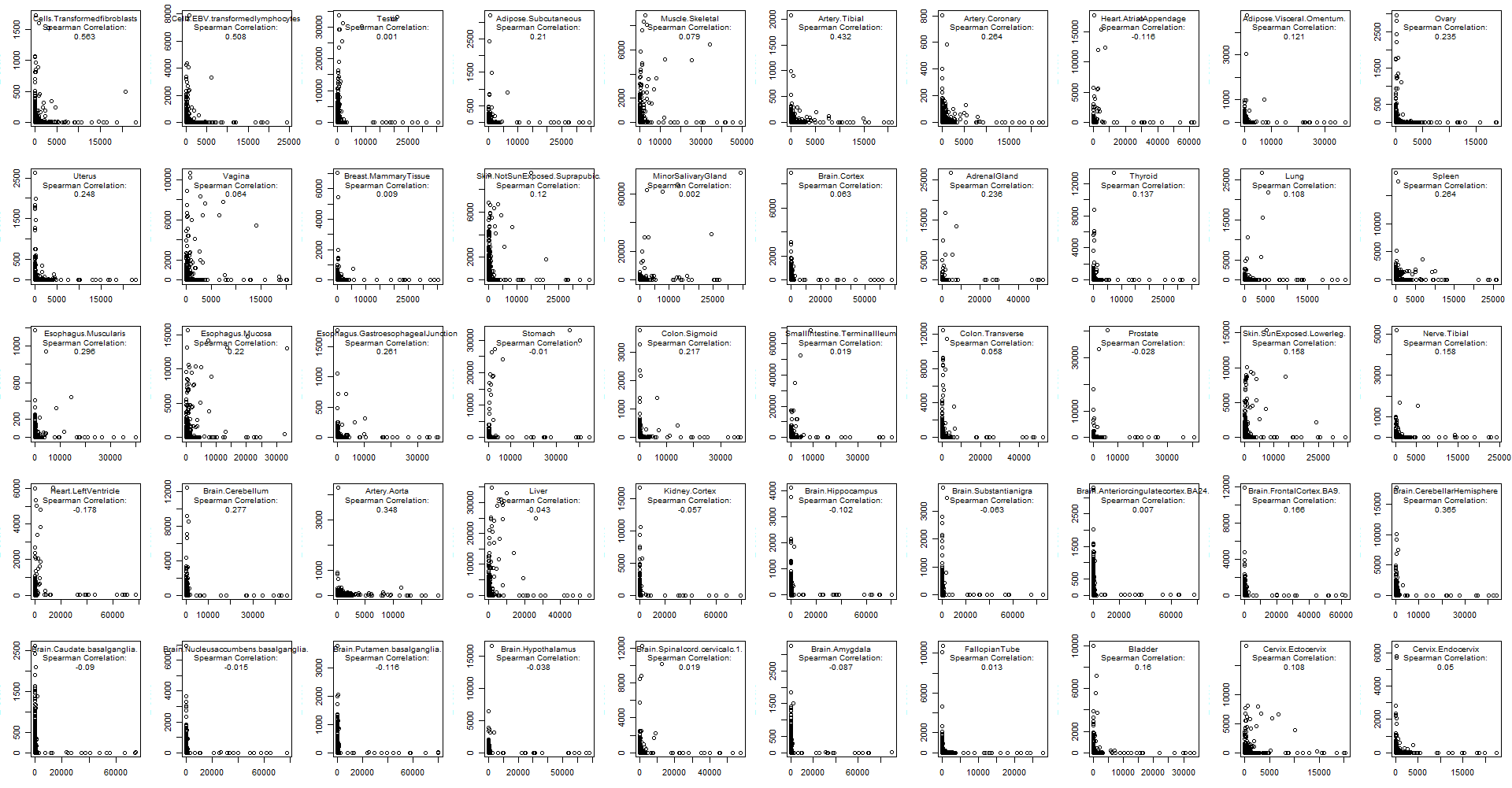

Figure S8: The scatter plots and correlation of original expression score and tissue specific scores at gene levels. The horizontal axis denotes the TPM score and vertical axis denotes the robust z-scores.

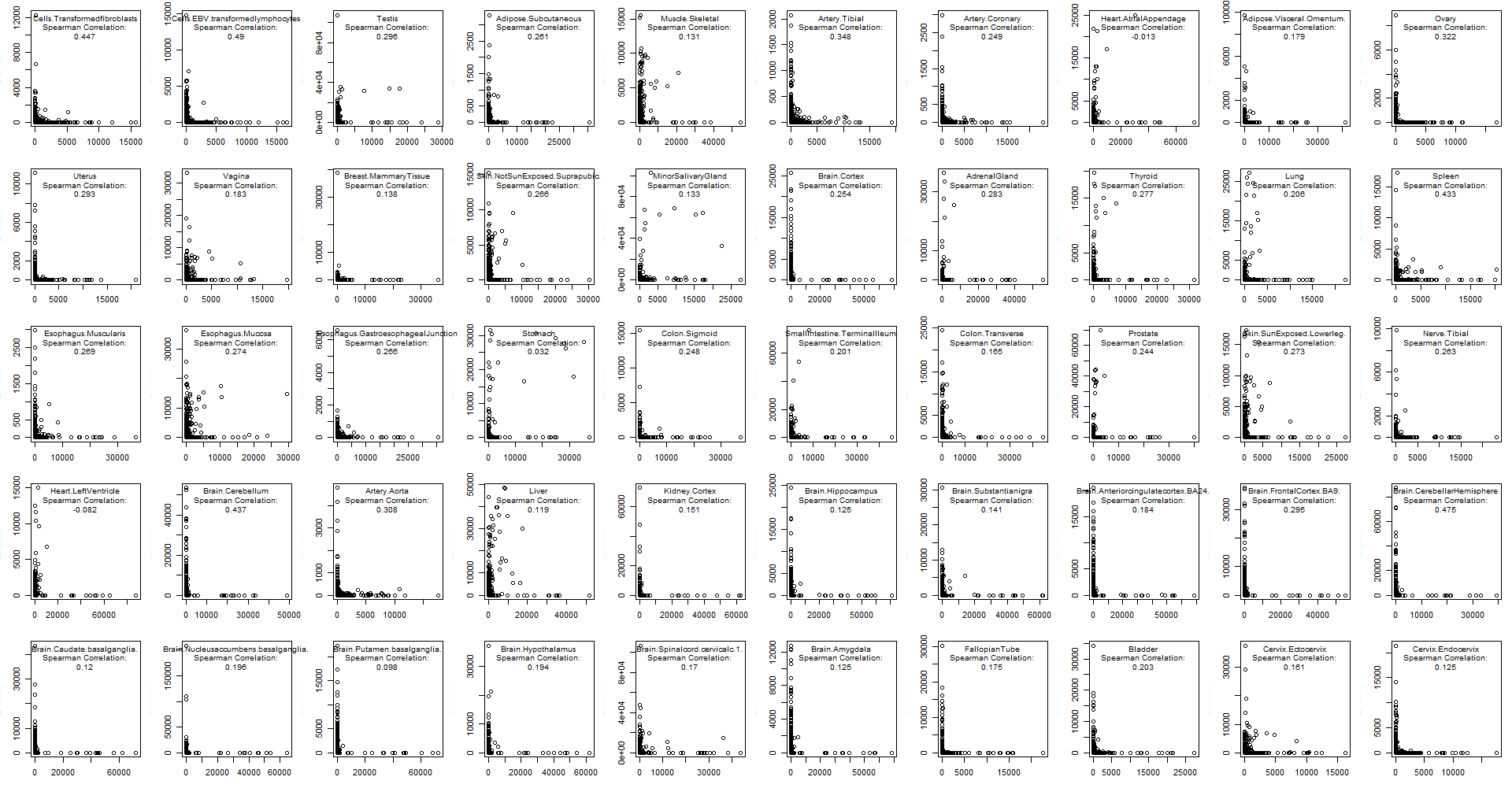

Figure S9: The scatter plots and correlation of original expression score and tissue specific expression scores at transcript levels. The horizontal axis denotes the TPM score and vertical axis denotes the robust z-scores.

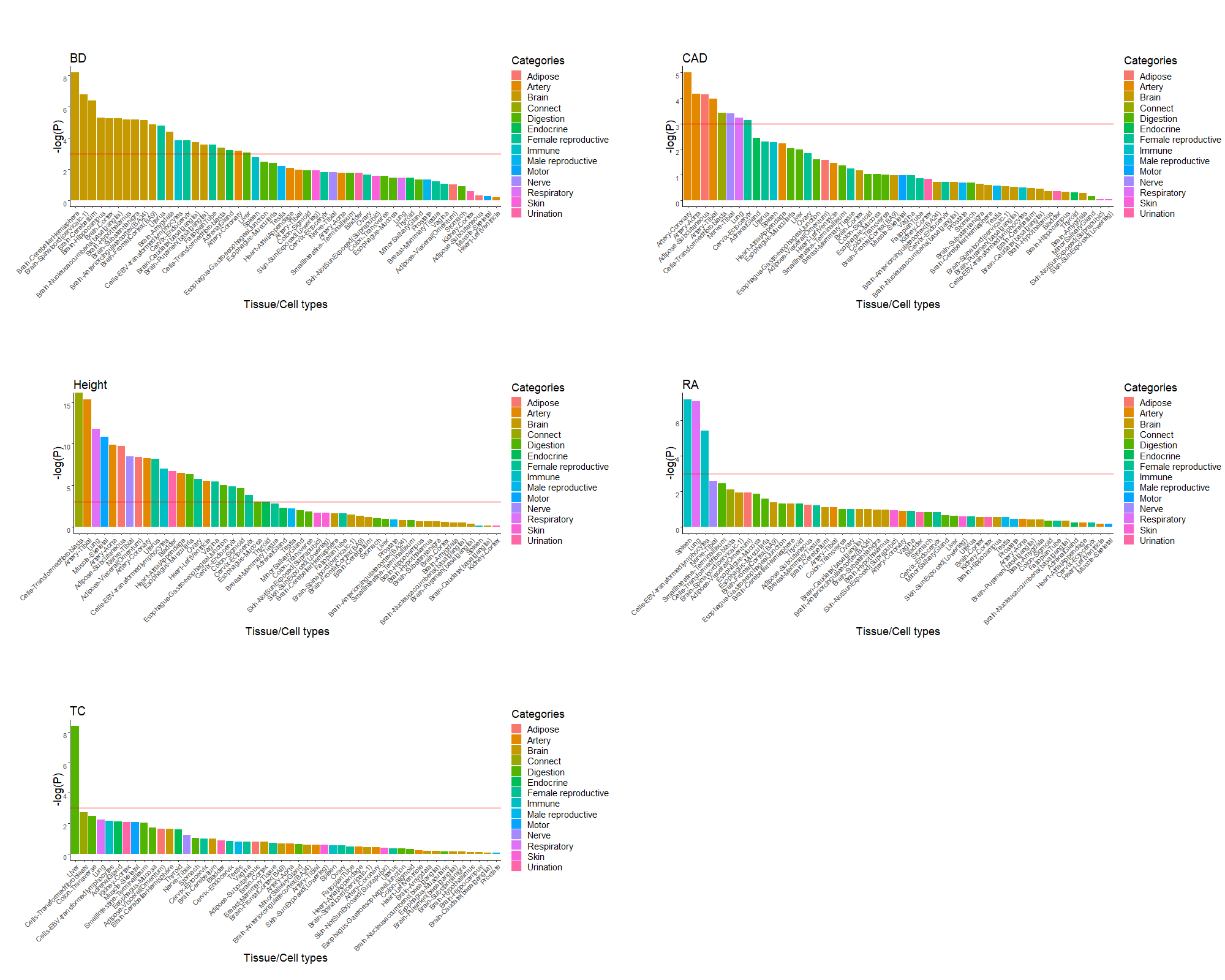

Figure S10: Prioritized driver tissues of five complex diseases/traits Note: on the *y* axis, the negative log_10_-transformed enrichment p-value of each tissue is plotted as bars. On the *x* axis, the 50 tissues are classified into 15 groups based on anatomy, which are filled with different colors as the legend shows. The tissues are sorted by the corresponding value on *y* axis in descending order. The p-values are calculated by hypergeometric distribution test for enrichment and the proposed robust z-score adopts a p-value cutoff 10^-6^. The tissue family-wise error rate is 0.05 and we adopt the standard Bonferroni correction for counteracting the problem of multiple comparison. And the red horizontal line denotes the log_10_-transformed p-value cutoff by 0.05/50=0.001. In the quality control of excluding extremely lowly expressed genes, gene or transcripts having TPM <0.05 in 40 or more tissues are excluded.

**Supplementary Tables**

Table 1: Statistical power of the robust z-score approach and standard z-score approach

| Number selectively expressed tissues | Expression value Deviated from mean (SD) | Power of the robust z-score(%) | Power of the conventional z-score(%) | Number selectively expressed tissues | Expression value Deviated from mean (SD) | Power of the robust z-score (%) | Power of the conventional z-score(%) |
| --- | --- | --- | --- | --- | --- | --- | --- |
| 1 | 1 | 100.0 | 100.0 | 1 | -1 | 100.0 | 100.0 |
|  | 2 | 100.0 | 100.0 |  | -2 | 100.0 | 100.0 |
|  | 3 | 100.0 | 100.0 |  | -3 | 100.0 | 100.0 |
| 3 | 1 | 99.9 | 78.8 | 3 | -1 | 100.0 | 78.7 |
|  | 2 | 100.0 | 99.7 |  | -2 | 100.0 | 99.7 |
|  | 3 | 100.0 | 100.0 |  | -3 | 100.0 | 100.0 |
| 6 | 1 | 80.4 | 0.5 | 6 | -1 | 80.3 | 0.5 |
|  | 2 | 99.9 | 0.0 |  | -2 | 99.9 | 0.0 |
|  | 3 | 100.0 | 0.0 |  | -3 | 100.0 | 0.0 |

Note: Fifty expression values were generated under a normal distribution with mean 10 and standard deviation 10, *N*(10, 10). Each power is estimated by 100,000 simulated datasets.

Table S2: Sample sizes of different tissues

| Tissue/CellType | Size |
| --- | --- |
| Adipose-Subcutaneous | 442 |
| Adipose-Visceral(Omentum) | 355 |
| AdrenalGland | 190 |
| Artery-Aorta | 299 |
| Artery-Coronary | 173 |
| Artery-Tibial | 441 |
| Bladder | 11 |
| Brain-Amygdala | 100 |
| Brain-Anteriorcingulatecortex(BA24) | 121 |
| Brain-Caudate(basalganglia) | 160 |
| Brain-CerebellarHemisphere | 136 |
| Brain-Cerebellum | 173 |
| Brain-Cortex | 158 |
| Brain-FrontalCortex(BA9) | 129 |
| Brain-Hippocampus | 123 |
| Brain-Hypothalamus | 121 |
| Brain-Nucleusaccumbens(basalganglia) | 147 |
| Brain-Putamen(basalganglia) | 124 |
| Brain-Spinalcord(cervicalc-1) | 91 |
| Brain-Substantianigra | 88 |
| Breast-MammaryTissue | 290 |
| Cells-EBV-transformedlymphocytes | 130 |
| Cells-Transformedfibroblasts | 343 |
| Cervix-Ectocervix | 6 |
| Cervix-Endocervix | 5 |
| Colon-Sigmoid | 233 |
| Colon-Transverse | 274 |
| Esophagus-GastroesophagealJunction | 244 |
| Esophagus-Mucosa | 407 |
| Esophagus-Muscularis | 370 |
| FallopianTube | 7 |
| Heart-AtrialAppendage | 297 |
| Heart-LeftVentricle | 303 |
| Kidney-Cortex | 45 |
| Liver | 175 |
| Lung | 427 |
| MinorSalivaryGland | 97 |
| Muscle-Skeletal | 564 |
| Nerve-Tibial | 414 |
| Ovary | 133 |
| Pancreas | 248 |
| Pituitary | 183 |
| Prostate | 152 |
| Skin-NotSunExposed(Suprapubic) | 387 |
| Skin-SunExposed(Lowerleg) | 473 |
| SmallIntestine-TerminalIleum | 137 |
| Spleen | 162 |
| Stomach | 262 |
| Testis | 259 |
| Thyroid | 446 |
| Uterus | 111 |
| Vagina | 115 |
| WholeBlood | 407 |

Table S3: Number of significantly expressed genes according to the robust z-scores at transcript

| Tissue/Cell Type | By gene expression | By transcript expression | Overlapped significant genes | Uniquely significant by gene expression | Uniquely significant by transcript expression |
| --- | --- | --- | --- | --- | --- |
| Adipose-Subcutaneous | 1864 | 3185 | 1507 | 357 | 1678 |
| Adipose-Visceral(Omentum) | 2107 | 3358 | 1675 | 432 | 1683 |
| AdrenalGland | 1539 | 3226 | 1245 | 294 | 1981 |
| Artery-Aorta | 1899 | 3522 | 1504 | 395 | 2018 |
| Artery-Coronary | 1842 | 2977 | 1437 | 405 | 1540 |
| Artery-Tibial | 1611 | 3701 | 1302 | 309 | 2399 |
| Bladder | 3413 | 8278 | 2427 | 986 | 5851 |
| Brain-Amygdala | 2348 | 4983 | 1867 | 481 | 3116 |
| Brain-Anteriorcingulatecortex(BA24) | 2685 | 5813 | 2126 | 559 | 3687 |
| Brain-Caudate(basalganglia) | 2682 | 5584 | 2110 | 572 | 3474 |
| Brain-CerebellarHemisphere | 4910 | 12133 | 3742 | 1168 | 8391 |
| Brain-Cerebellum | 5073 | 12302 | 3788 | 1285 | 8514 |
| Brain-Cortex | 3104 | 6519 | 2425 | 679 | 4094 |
| Brain-FrontalCortex(BA9) | 3287 | 7205 | 2629 | 658 | 4576 |
| Brain-Hippocampus | 2525 | 5365 | 2037 | 488 | 3328 |
| Brain-Hypothalamus | 3113 | 6413 | 2469 | 644 | 3944 |
| Brain-Nucleusaccumbens(basalganglia) | 2972 | 6346 | 2330 | 642 | 4016 |
| Brain-Putamen(basalganglia) | 2221 | 4701 | 1781 | 440 | 2920 |
| Brain-Spinalcord(cervicalc-1) | 2834 | 5717 | 2199 | 635 | 3518 |
| Brain-Substantianigra | 2343 | 4824 | 1885 | 458 | 2939 |
| Breast-MammaryTissue | 2543 | 3834 | 1893 | 650 | 1941 |
| Cells-EBV-transformedlymphocytes | 3765 | 8080 | 2876 | 889 | 5204 |
| Cells-Transformedfibroblasts | 2278 | 6923 | 1801 | 477 | 5122 |
| Cervix-Ectocervix | 3717 | 8457 | 2560 | 1157 | 5897 |
| Cervix-Endocervix | 4174 | 9917 | 2780 | 1394 | 7137 |
| Colon-Sigmoid | 1592 | 2683 | 1314 | 278 | 1369 |
| Colon-Transverse | 2429 | 3727 | 1929 | 500 | 1798 |
| Esophagus-GastroesophagealJunction | 1136 | 2037 | 971 | 165 | 1066 |
| Esophagus-Mucosa | 2459 | 4173 | 1958 | 501 | 2215 |
| Esophagus-Muscularis | 1064 | 2111 | 921 | 143 | 1190 |
| FallopianTube | 4491 | 10142 | 3062 | 1429 | 7080 |
| Heart-AtrialAppendage | 1027 | 2322 | 840 | 187 | 1482 |
| Heart-LeftVentricle | 675 | 1785 | 549 | 126 | 1236 |
| Kidney-Cortex | 2328 | 4378 | 1773 | 555 | 2605 |
| Liver | 1922 | 3698 | 1536 | 386 | 2162 |
| Lung | 4107 | 6807 | 3162 | 945 | 3645 |
| MinorSalivaryGland | 2825 | 4597 | 2125 | 700 | 2472 |
| Muscle-Skeletal | 1170 | 3365 | 973 | 197 | 2392 |
| Nerve-Tibial | 2841 | 5033 | 2056 | 785 | 2977 |
| Ovary | 2945 | 4360 | 1811 | 1134 | 2549 |
| Prostate | 3181 | 5430 | 2449 | 732 | 2981 |
| Skin-NotSunExposed(Suprapubic) | 2760 | 4811 | 2184 | 576 | 2627 |
| Skin-SunExposed(Lowerleg) | 2656 | 4829 | 2140 | 516 | 2689 |
| SmallIntestine-TerminalIleum | 3394 | 5489 | 2686 | 708 | 2803 |
| Spleen | 3917 | 6456 | 2938 | 979 | 3518 |
| Stomach | 2002 | 2790 | 1487 | 515 | 1303 |
| Testis | 12520 | 26086 | 8911 | 3609 | 17175 |
| Thyroid | 3382 | 5766 | 2387 | 995 | 3379 |
| Uterus | 2147 | 3806 | 1510 | 637 | 2296 |
| Vagina | 2920 | 4688 | 2219 | 701 | 2469 |

Note: The Bonferroni correction was used to control family-wise error rate of 0.05 on the entire genome for declaring significantly expressed genes. When using z-score at transcripts, the cutoff for a gene was further divided by the number of transcripts according to the Bonferroni correction.

Table S4: GWAS summary results information of five complex diseases/traits

| **Trait name (short)** | **Sample size** | **Marker counts** | **Web link** |
| --- | --- | --- | --- |
| Bipolar disorder | 41,653 | 8,949,599 | <https://www.med.unc.edu/pgc/results-and-downloads> |
| Rheumatoid arthritis (RA) | 103,638 | 9,739,303 | <https://grasp.nhlbi.nih.gov/downloads/ResultsOctober2016/Okada/RA_GWASmeta_TransEthnic_v2.txt.gz> |
| Coronary artery disease | 184,305 | 9,446,655 | <http://www.cardiogramplusc4d.org/media/cardiogramplusc4d-consortium/data-downloads/cad.additive.Oct2015.pub.zip> |
| Total cholesterol (TC) | 188,577 | 2,446,981 | <https://grasp.nhlbi.nih.gov/downloads/ResultsOctober2016/Global_Lipids_Genetics_Consortium/jointGwasMc_TC.txt.gz> |
| Height | 253,288 | 2,547,478 | <https://grasp.nhlbi.nih.gov/downloads/ResultsOctober2016/Wood/GIANT_HEIGHT_Wood_et_al_2014_publicrelease_HapMapCeuFreq.txt.gz> |
